# Quantifying genetic regulatory variation in human populations improves transcriptome analysis in rare disease patients

**DOI:** 10.1101/632794

**Authors:** Pejman Mohammadi, Stephane E. Castel, Beryl B. Cummings, Jonah Einson, Christina Sousa, Paul Hoffman, Sandra Donkervoort, Payam Mohassel, Reghan Foley, Heather E. Wheeler, Hae Kyung Im, Carsten G. Bonnemann, Daniel G. MacArthur, Tuuli Lappalainen

## Abstract

Transcriptome data holds substantial promise for better interpretation of rare genetic variants in basic research and clinical settings. Here, we introduce ANalysis of Expression VAriation (ANEVA) to quantify genetic variation in gene dosage from allelic expression (AE) data in a population. Application to GTEx data showed that this variance estimate is robust across datasets and is correlated with selective constraint in a gene. We next used ANEVA variance estimates in a Dosage Outlier Test (ANEVA-DOT) to identify genes in an individual that are affected by a rare regulatory variant with an unusually strong effect. Applying ANEVA-DOT to AE data form 70 Mendelian muscular disease patients showed high accuracy in detecting genes with pathogenic variants in previously resolved cases, and lead to one confirmed and several potential new diagnoses in cases previously unresolved. Using our reference estimates from GTEx data, ANEVA-DOT can be readily incorporated in rare disease diagnostic pipelines to better utilize RNA-seq data.

**One Sentence Summary:** New statistical framework for modelling allelic expression characterizes genetic regulatory variation in populations and informs diagnosis in rare disease patients

---

Large reference databases of human exomes and genomes have enabled characterization of genomic variation in human populations (*1–3*). These data have been used to summarize genic intolerance to damaging variants (*1, 4, 5*), which has been essential for prioritizing rare and *de novo* coding variants to achieve genetic diagnosis of Mendelian disease currently for 25-50% patients (*6, 7*). Yet, a large number of patients remain undiagnosed after exome and/or whole genome sequencing. Despite remarkable advances in DNA sequencing, the search for rare disease-causing variants outside the coding sequence has been hindered by the difficulty of interpreting rare regulatory variants and identifying their target genes.

Integration of genome and transcriptome sequencing data has provided promising early results of improved diagnosis via better detection of rare variants with functional effects (*6, 8, 9*). However, this analysis remains limited to a small fraction of variants that induce clear alterations in the transcriptome, such as stop gains and splice defects, and is often highly laborious involving manual inspection of RNA-seq reads by experts in patients and reference transcriptome cohorts. Furthermore, the dynamic nature of the transcriptome with changes driven by environment, disease state, and technical variation has made it challenging to quantify when a rare genetic effect that is beyond the normal range of population variation. A promising data type that can capture genetic effects is allelic expression (AE) which measures the relative expression of the paternal and maternal haplotype of a gene in an individual. Departure from equal AE, allelic imbalance, has unique sensitivity in capturing genetic effects in *cis* as it is largely unaffected by environmental and technical factors with a reported heritability of 85% (*10*). Allelic imbalance can detect signatures of rare regulatory variation (*6, 11–13*), and thus has potential for a comprehensive quantification of genetic regulatory variants across the frequency spectrum (*14*). However, a quantitative framework for interpreting this unique data type to identify pathogenic rare variants has been lacking. In this work, we use a mechanistic model of *cis*-regulatory variation to quantify genetic regulatory variation in populations. Specifically, for each gene we estimate the expected population variance in its dosage that is due to genetic differences among individuals. We refer to this quantity as V^G^. We show that V^G^ can be robustly estimated from publicly available transcriptome data and can be used as a population reference for identification of genes affected by pathogenic variants in patients with RNAseq data.

The effect sizes of *cis*-regulatory variants can be quantified using allelic fold change (aFC; (*15*)). The analytical link between aFC and gene dosage would allow us to calculate V^G^ if the effect sizes and haplotype frequencies of all common and rare regulatory variants were known (**Eq. 1–7**). Since this is in practice never the case, we instead use AE data to infer their overall distribution without having to identify individual regulatory variants explicitly. Across individuals, AE data represents a series of comparisons of net expression effects of all variants on two random haplotypes at a time. In a population sample, this forms a wide range of patterns depending on the properties of regulatory variants present in the population and their linkage with aeSNP that is the SNP used to measure allelic expression (**Figure 1A-D**), which has been a major complication for applications of AE data (*13, 14*). We extended the mechanistic model of aFC to derive a generative model for population AE data under a realistic scenario in which a given gene is regulated by several regulatory variants only some of which are known (or statistically identifiable). We show that under this assumption population AE data is described by a constrained mixture of Binomial-Logit-Normal (BLN) probability distribution functions (**Eq. 8–19**). We fit this model to population AE data (**Eq. 20–28, Figure 1E-F, Fig S1A-B**), and use the maximum likelihood parameters to estimate V^G^ indirectly (**Eq. 29–30, Figure 1G**). We refer to this method of estimating V^G^ as ANalysis of Expression Variation (ANEVA). We evaluated the performance of ANEVA by simulating AE data with 1-15 regulatory variants per gene with varying allele frequencies, with highly accurate V^G^ estimates (R^2^=92%, **Fig S1**). Thus, ANEVA allows translation of allelic expression read count data into biologically interpretable estimates of genetic variation in gene expression in populations.

**Figure 1:**
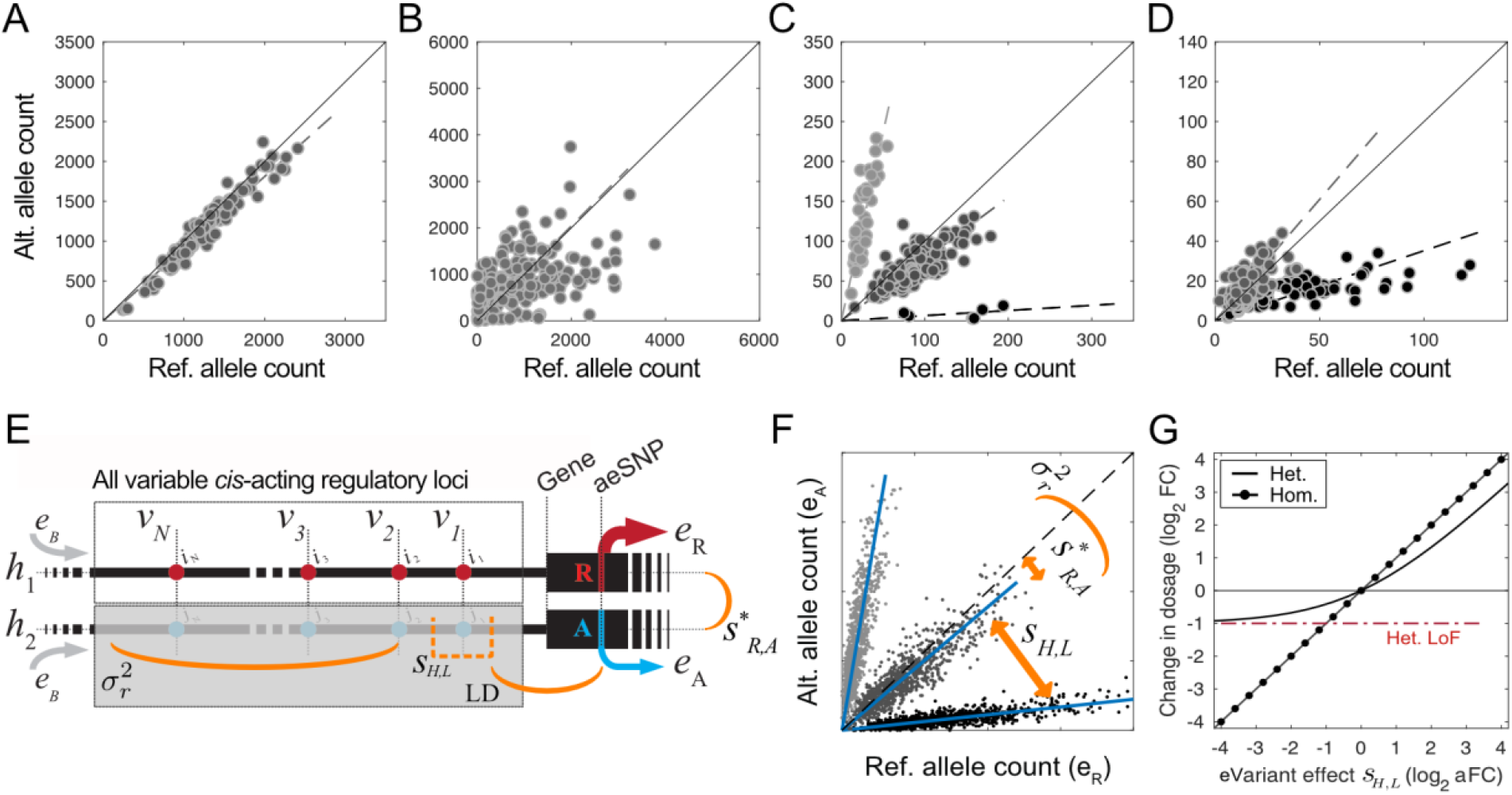
*Cis*-regulatory variation, allelic expression, and ANEVA. A-D) Examples of allelic expression patterns in a population sample for four genes. Each dot represents an individual and each panel is produced from a single aeSNP. The gene in (A) has little *cis*-regulatory variation since all individuals have very similar expression levels of both haplotypes, whereas in (B) there is scattered variation without clear clusters that would represent a single strong variant. The patterns in (C) and (D) show distinct clusters that are driven by different haplotype combinations of a common, strong regulatory variant and the aeSNP. In (D) one of the clusters is eliminated due to strong linkage disequilibrium. These examples illustrate how different AE patterns can be, highlighting the challenge of consistently modeling the underlying regulatory variants and their effect sizes and frequencies. E-F) Population model for AE data. AE data is modelled using one distinctly strong regulatory bi-allelic variant. If present, this variant is specified by its effect size, *s*_H,L_, and its LD with the aeSNP. Residual *cis*-regulatory variation is modelled as an infinite-allelic regulatory variant summarized by variance term 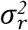. Allelic expressions *e*_R_ and *e*_A_ are measured at a heterozygous aeSNP with reference (R) and alternative (A), and *s*\*_R,A_ is the effect size of the reference allele alignment bias for the aeSNP. Haplotypes *h*_1_ and *h*_2_, shared basal expression level *e*_B_, and *N cis*-regulatory variant sites *v*_1_…*v_N_*, are component of our complete formal model of *cis*-regulatory variation which were used for derivation of the population model for AE data and for simulation studies. G) The relationship of effect size of a regulatory variant measured as allelic fold change (x-axis) and total gene expression dosage (y-axis).

We applied ANEVA to 10,361 RNA-seq samples spanning 48 tissues and 620 individuals with WGS data from the GTEx v7 data set (*16, 17*). We estimated V^G^ over a total of 2,150,010 autosomal aeSNPs with at least 30 reads in 6 individuals, separately in each tissue. We derived gene-level estimates of V^G^ as weighted harmonic mean of SNP-level estimates for a median of 4,962 genes per tissue, and a total of 14,084 genes are present in at least one tissue (**Figure 2A-B, Table S1**). To ensure that our AE-derived estimates reflect classical eQTL mapping of regulatory variation, we calculated V^G^ estimates using eQTL effect size and allele frequencies for each gene in each tissue, with consistent results (median rho = 0.75; **Figure 2C, Fig S2, Table S2**). We also benchmarked ANEVA estimates against estimates of gene expression *cis*-heritability. For each gene in GTEx whole blood data, we calculated the ratio of AE and eQTL derived V^G^ estimates from ANEVA to the total variance of gene expression (V^T^), and a standard local heritability estimate calculated by Bayesian Sparse Linear Mixed Model (BSLMM; (*18*)). Comparing these to BSLMM estimates from the larger DGN dataset (922 donors versus 369 in GTEx whole blood) as gold standard, we found that GTEx whole blood estimates from ANEVA are well correlated and better powered than BSLMM applied to GTEx (**Figure 2D; Fig S3**). The fact that ANEVA estimates based on a single eQTL per gene yield a higher correlation than AE-based estimates may be due to BSLMM mostly capturing common variant effects at the current sample sizes. Since AE-based ANEVA V^G^ estimates are better applicable to AE-based outlier detection detailed below, subsequent analyses were based on them (see **Supplementary text 1**).

**Figure 2:**
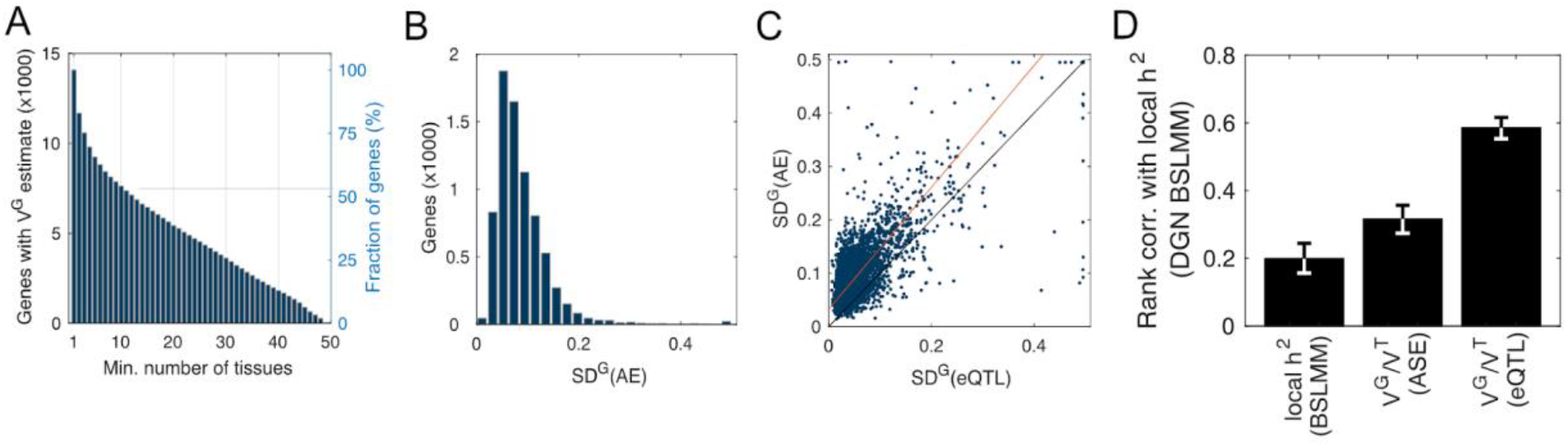
Estimates of genetic regulatory variation in GTEx. A) Number of genes that have V^G^ estimates across 1 to 49 GTEX tissues; B-C) Distribution of SD^G^, square root of V^G^, for 7556 genes available in GTEx adipose subcutaneous tissue (B), and its comparison to estimates derived from eQTL data (C). The red line is Deming regression fit (see **Fig S2** for additional details). D) Correlation of GTEx whole blood gene expression heritability estimates (h^2^) from ANEVA and local h^2^ from the traditional linear mixed model-based approach (BSLMM). Estimates derived by BSLMM from the much larger DGN data was used as reference. Our V^G^-based h^2^ estimates from GTEx data are well correlated to BSLMM estimates from DGN data (see **Supplementary text 1**). SD^G^ is capped at 0.5 for visualization in panels B and C.

Next, we analyzed how V^G^ varies between tissues and populations. We found that estimates of V^G^ between two tissues tend to be well correlated (median rho=0.57) and follow general tissue relatedness (**Figure 3A**). Furthermore, for a given gene V^G^ tends to be smaller in tissues were the gene is higher expressed (Wilcoxon signed rank test *p*<10^−300^; **Figure 3B**). We verified by down-sampling and analysis of eQTL-derived V^G^ estimates that this pattern was not an artifact of differences in read depth (**Fig S4**). These results indicate that genetic variation has a smaller absolute effect on gene dosage in tissues where the gene has higher expression, which could be a sign of increased dosage sensitivity and higher selective constraint in tissues where the gene has a more pronounced functional role. A possible example of this is *PAX8*, a transcription factor with high expression in thyroid, that has a haploinsufficient association with congenital hypothyroidism (*19*). While *PAX8* harbors a strong common eQTL introducing substantial dosage variation in other tissues, in thyroid this eQTL is not active (**Fig S5**). Finally, to analyze population differences in V^G^, we used ANEVA on AE data from three European subpopulations (FIN=89, GBR=86, TSI=92) and one African subpopulation (YRI=77) from GEUVADIS (*20*). V^G^ estimates were highly correlated with little signs of higher concordance between the European subpopulations. This suggests that the total amount of genetic dosage variation is a general property of a gene that is not highly affected by genetic differences between human populations (**Fig S6**). This suggests that approaches that aggregate genetic effects at the gene level may have promise in better applicability across populations than analyses of individual variants.

**Figure 3:**
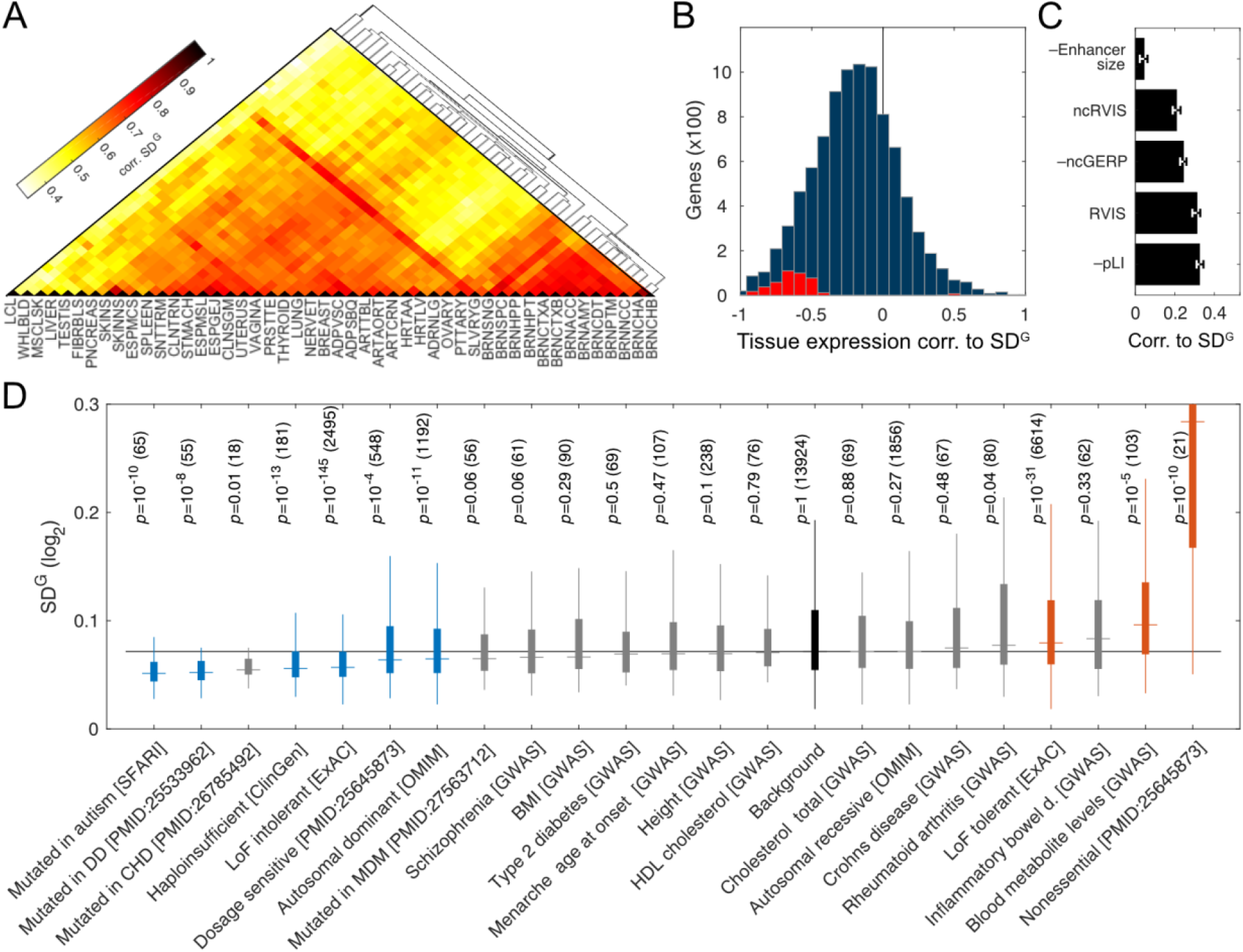
Biological sources of regulatory variation between genes. A) Correlation of genetic regulatory variation across GTEx tissues B) Correlation between median expression in a tissue and V^G^ for 9,158 genes with V^G^ estimates in at least five tissues. Overall distribution is biased to towards negative values (Med. rho −0.20). Genes individually showing significant correlation are shown in red (5% FDR). C) Correlation of 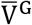, average V^G^ across tissues, with estimates of enhancer size, coding constraint (RVIS, pLI), and noncoding constraint (ncRVIS) and conservation (ncGERP) in UTRs and promoters. D) Distribution boxplots of 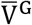 values for different gene sets (DD: Developmental disorder, CHD: Congenital heart disease, MDM: Congenital Muscular dystrophies and myopathies; **Table S3**), with nominal p-values from ranksum test compared to the background of all genes, followed by the number of genes in each set in parentheses. Gene sets with p-value ≤ 0.01 are colored. Boxes span the middle 50% values, and the whiskers span ±1.5 IQR from first and the third quartile. SD^G^ is the square root of V^G^.

In order to understand the biological processes that lead to differences in the amount of genetic regulatory variation between genes, we correlated our V^G^ estimates to statistics of gene regulation and constraint. First we aggregated tissue-specific estimates of V^G^ for each gene using harmonic mean weighted by gene expression in each tissue (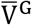, **Table S1**). We observed a minimal correlation between 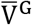 and the estimated enhancer size (**Figure 3B**; (*21*)), which suggests that the size of the mutational target and probably also the background mutation rate play a minor role in the total amount of genetic regulatory variation. Genes with high purifying selection for coding gene disrupting variants, or noncoding variants in the promoter or UTR regions, were also depleted of genetic regulatory variation (**Figure 3C**), as observed before by analyzing the depletion of eQTLs (*1*). Next, we explored the distribution of 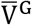 in a range of gene sets (**Table S3**). We found that genes implicated in rare diseases by exome sequencing studies are depleted of genetic regulatory variation, and those tolerant of loss of function mutations tend to also tolerate more genetic regulatory variation (**Figure 3D**). This points to dosage sensitivity that is captured by both exome sequencing studies and our 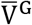 analysis. GWAS genes for different traits and diseases showed little deviation from the background but had patterns consistent with genetic architecture of these traits, with GWAS genes for schizophrenia risk having the lowest and blood metabolite genes the highest V^G^. While the exact contribution of purifying selection to gene expression variation is difficult to quantify (*14, 22*), these results indicate that the amount of genetic regulation variation measured as V^G^ can be a proxy for regulatory constraint of different genes, complementing estimates derived from coding variation. This provides insights into genetic architecture of human traits.

In addition to the general biological insights provided by the V^G^ estimates, they have a direct practical application in identifying population outliers. To this end, we developed ANEVA Dosage Outlier Test (ANEVA-DOT), that utilize ANEVA’s V^G^ estimates to identify genes where an individual appears to carry a genetic variant affecting gene dosage that has an unusually strong effect relative to what is expected in general population (**Eq. 31–42; Figure 4A**). We hypothesized that in a patient such dosage outlier genes are more likely to be pathogenic. Briefly, ANEVA-DOT compares an individual’s allelic counts for each gene to the general population variation represented by V^G^, and accounting for a number of additional technical and biological sources of variation (**Figure 4A**). We ensured that the test is well calibrated by extensive simulations (**Fig S7**).

**Figure 4:**
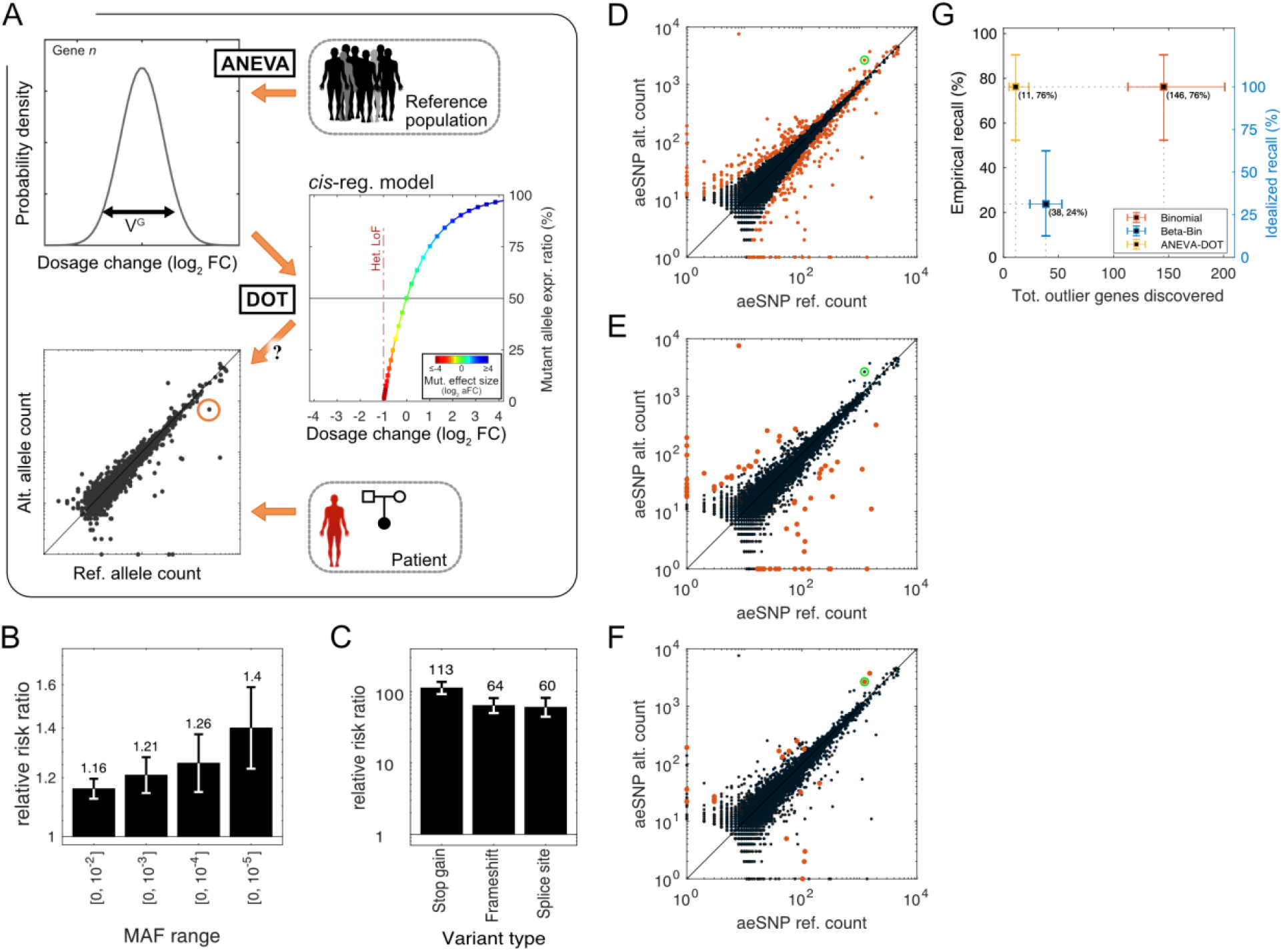
Regulatory outliers detected with the ANEVA Dosage Outlier Test. A) Illustration of the ANEVA-DOT method. It uses the population V^G^ estimate of variance in each gene and the model of *cis*-regulatory genetic effect to estimate the null distribution of allelic imbalance. An individual’s allelic counts for that gene are compared to this null, accounting for sampling noise, sequencing noise, reference bias and the variant haplotype. B) Progressive enrichment of all rare variants in ANEVA-DOT genes as a function of allele frequency. (C) Enrichment of rare putative gene-disrupting variants (MAF<1%) in ANEVA-DOT genes. D-F) An example of allelic expression across genes in one previously diagnosed (green circle) muscle dystrophy patient, with allelic expression outliers marked in red from binomial (*n*=407; D), beta-binomial (*n*=66; E), and ANEVA-DOT (N=19; F) tests. G) Fraction of true causal genes identified (recall) versus the number reported outliers for previously diagnosed patients where AE would be expected. Empirical recall (left) is calculated using all patients including cases where identifying the gene is essentially impossible (e.g. causal gene is not expressed), while idealized recall (right) is limited to those cases where the causal gene is present in the AE data and there is allelic imbalance.

To test ANEVA-DOT in the general population, we applied it to 466 skeletal muscle samples from GTEx individuals. A median of 3,390 genes were tested in each sample resulting in a median of 10 genes (90% range: [3, 22]) identified as outliers by ANEVA-DOT (5% FDR; hereafter ANEVA-DOT genes). As a quality filter, we discarded for further analysis 113 out of 5848 tested genes that appeared as outliers in >1% of the individuals likely due to noisy V^G^ estimates or low-quality AE data (**Fig S8, Table S4**). After this step each sample retained a median of 4.5 ANEVA-DOT genes (90% range: [1, 14]; **Figure 4B**). Looking within a 10kb window upstream of the TSS and the gene body of ANEVA-DOT genes, we found that they are enriched for rare heterozygous variants (**Fig 4B**). As a positive control, we verified that this enrichment is particularly strong for rare putative gene disrupting ones where a strong effect on gene expression levels (via nonsense-mediated decay) is expected (**Figure 4B-C**). This confirms that ANEVA-DOT picks up rare genetic effects on gene dosage.

Next, we evaluated how sensitive ANEVA-DOT is to differences in the reference population where V^G^ is calculated, since perfectly matching reference data may not always be available. First, using the GEUVADIS data described above, we looked for ANEVA-DOT genes in each of the 86 GBR European individuals using V^G^ estimates derived from two European (FIN and TSI), and one African (YRI) populations. An average of 74% (69%–78%) of ANEVA-DOT genes identified using one of the reference population are confirmed by a second population with YRI performing equally to the European populations (**Fig S9**). This supports the previous observation that analyses based on gene-level V^G^ have a reasonably good applicability across populations. Next, we looked at ANEVA-DOT genes identified in GTEx skeletal muscle samples that can be identified with V^G^ reference from fibroblast, adipose subcutaneous, skin sun-exposed, and whole blood instead of muscle. Fibroblast had the best replication rate with 23.3% of ANEVA-DOT genes found, with only 12.3% in whole blood. This is due to differences in both V^G^ and gene expression and indicates that a matching reference tissue is important (**Fig S10**).

To test ANEVA-DOT’s performance in diagnosis of rare disease patients, we applied it to AE data from 70 rare Mendelian muscle dystrophy and myopathy (MDM) patients, with V^G^ reference from GTEx skeletal muscle (**Fig S11–S17, Table S5**). Out of the 65 patients with high quality AE data, 32 have a previous diagnosis of which 21 are expected to lead to allelic imbalance (*6*), which were used as positive controls to benchmark ANEVA-DOT against binomial and beta-binomial tests of allelic imbalance (**Figure 4D-F**). At 5% FDR, binomial test identified a median of 146 genes with allelic imbalance signal per individual. These included the previously diagnosed disease gene in 76% (*n*=16) of the individuals. In the remaining 24% of the cases, AE data was not available for the disease gene (*n*=3) or no allelic imbalance was observed (*n*=2; **Fig S12**). These represented the fraction of cases that are essentially not amenable to any AE-based test. ANEVA-DOT identified a median of 11 outlier genes per individual (out of a median of 2190 with sufficient data), including genes where a subtle allelic imbalance in absolute terms is still an outlier due to very low general population variance (**Figure 4F**). The previously diagnosed gene was identified among the ANEVA-DOT genes in all 76% (*n*=16) of the individuals where there was a detectable allelic imbalance present (**Fig S11–S12**). Notably, in 69% (*n*=11) of these cases the causal gene was among the top-five most significant tested genes by p-value (**Table S5**). In contrast, a simple beta-binomial test captured the diagnosed gene only in 24% (*n*=5) of the cases where the disease gene had a considerably strong allelic imbalance. Thus, ANEVA-DOT presented a considerable improvement in precision as compared to binomial test while maintaining sensitivity in contrast to a beta-binomial test (**Figure 4G**). Furthermore, using data from 14 patients with previous diagnosis that are not expected to show allelic imbalance as negative controls, we confirmed that ANEVA-DOT leads to significantly lower false positive rate that the other two tests (**Fig S12–S14, See Supplementary text 2**).

In the remaining 33 patient samples without a genetic diagnosis with WES and/or WGS or RNA-seq analysis with earlier pipelines (*6*), we found a median of 9 ANEVA-DOT genes per sample (in total 349 genes), and in 12 patients this included at least one gene (in total 17 genes) previously implicated in neuromuscular disorders (**Fig S15–S16**; (*6, 23*)). By the time of writing this report, one new diagnosis from ANEVA-DOT was confirmed for a patient in the previously published Cummings et al. (*6*) cohort. Patient N10 from Cummings et al, with limb-girdle muscular dystrophy-like phenotype had gone through prior gene panel, WES, WGS and RNA-seq analysis. ANEVA-DOT identified 13 outlier genes in this patient, with *DES* being the most significant one by p-value and the only known Mendelian muscle disease gene on the list. Follow-up evaluation of the RNA-seq data in the gene identified a pseudoexon insertion caused by an intronic splice site creating variant. This splice aberration was missed due to the relatively low number of reads in support but has now been confirmed by RT-PCR. The variant is found *in trans* with a previously reported pathogenic missense variant in *DES* that had not been identified as a diagnosis due to the lack of a second variant. This demonstrates the utility of ANEVA-DOT for detecting previously undiscovered compound heterozygotes (**Fig S18**). Additionally, ANEVA-DOT prioritized strong candidates in six cases, and identified possible candidates in 11 others (**See Supplementary text 3**). The strong candidates that ANEVA-DOT followed by WGS analysis is currently able to pinpoint are gene or splice disrupting variants, since the functional role of these variants, with the support of ANEVA-DOT, is easier to assign compared to rare regulatory variant candidates. Overall, we expect up to 10.5 of the 17 known MDM, and 18.8 of all 349 identified ANEVA-DOT genes in the 33 undiagnosed patients to be true disrupted causative genes (**See Supplementary text 2**).

Altogether, these results show that ANEVA-DOT is a valuable method for identifying genes where an individual is likely to have a rare heterozygous variant with a strong impact on gene expression, and that it has a high specificity and sensitivity in identifying disease-contributing genes when the causal variant affects gene dosage. By design, ANEVA-DOT does not rely on identifying which variant underlies the dosage outlier effect, since this is often unknown for rare regulatory variants, but genetic analysis can be applied after prioritizing genes by ANEVA-DOT. ANEVA-DOT is implemented in R, and it runs in a few seconds per sample.

## Discussion

In this study, we have introduced a novel method, ANalysis of Expression VAriation (ANEVA), and its extension ANEVA Dosage Outlier Test (ANEVA-DOT) to quantify genetic variation in gene dosage in the general population, and to identify genes where a patient appears to carry a heterozygous variant with an unusually strong effect in gene expression. This provides a methodology that allows for finding genes affected by potentially pathogenic variants by comparing individual transcriptome samples to previously generated reference data, in a manner that avoids the caveats of technical and reverse causation noise in total gene expression analysis. Further strengths of these methods include measuring variation in biologically interpretable units of gene dosage, which will allow interpretation of regulatory and coding gene disrupting variants on the same scale. The novel statistical methods introduced here for modeling allelic expression (AE) data are applicable to other uses of this data type. ANEVA analyses show that the amount of genetic regulatory variation per gene in the general population is highly correlated to selective constraint and is lowered in rare disease genes indicating dosage sensitivity. We demonstrate that ANEVA-DOT is a fast and powerful approach for finding genes with likely disease effects, with the small numbers of outliers making further manual curation feasible in a clinical setting without compromising on sensitivity.

V^G^ estimates from GTEx and other external data sets can be used as a shared reference for ANEVA-DOT analysis of patient data as long as the tissue is matched. This is analogous of coding constraint metrics that are essential for prioritization of pathogenic coding variants, and we now provide a similar framework for variants affecting gene expression levels. Since this method captures transcriptome outcomes of genetic effects without having to identify rare regulatory variants themselves, this method is particularly advantageous for capturing rare genetic effects from poorly defined regulatory elements, but it will also detect for example nonsense variants triggering transcript decay. We note however, that identifying and verifying the specific noncoding mutation resulting in the AE changes is still challenging.

Despite these advantages, our methods have several limitations. The main caveat is that while AE data is uniquely sensitive in detecting genetic regulatory effects in *cis*, the data has non-random sparsity. Genes with few common coding variants due to small size or high coding constraint can easily lack sufficient data for ANEVA’s V^G^ estimation, and lowly expressed genes can have noisy estimates. These issues will, however, improve with increasingly large RNA-seq data sets, and potential integration of V^G^ values derived from AE and eQTL data. In ANEVA-DOT, only about half of the genes expressed in each individual have AE data that is needed to be able to test them, and sometimes aeSNPs are in a position where they do not capture, for example, an in-frame splice change. Finally, allelic imbalance is not informative of recessive effects without family analysis. Thus, similarly to other genetic diagnosis tools, ANEVA-DOT should be used together with other methods to jointly capture different types of rare variants underlying disease. We envision that in clinical genetics, when practically feasible, transcriptome data will become a powerful additional layer of data for interpreting the genome and its disease-contributing variants. Sophisticated tools such as ANEVA-DOT are important in bringing this to practice, and we anticipate interesting applications and extensions of this approach to different genetic architectures and study designs.

## Acknowledgements

We would like to thank the GTEx consortium and the members of its analysis group and the rare variant working group for assistance in data processing and feedback on this work. In particular, we would like to thank Francois Aguet for help with AE data production, and Nicole Ferraro for processing of variant annotations, and Stephen Montgomery for insightful comments on the manuscript.

## Authors contributions

P.M. and T.L. designed the study. P.M. developed all statistical models, P.M., S.E.C., B.C., and J.E. analyzed the data, and P.M., C.S. and P.H. developed software tools. H.E.W., H.K.I., and D.G.M. provided data and materials. P.M. and T.L. wrote the paper with input from all the authors.

## Data and materials availability

Software packages for ANEVA, ANEVA-DOT, and BLN distribution functions are available in https://github.com/PejLab in individual repositories with corresponding names. Reference ANEVA estimates from **Table S1** and full ANEVA-DOT summary stats similar to **Table S4** for all GTEx tissues are available and will be updated in https://github.com/PejLab/Datasets. The GTEx v7 data is available in dbGap (phs000424.v7.p2), and the MDM cohort data is available in dbGap phs000655. Gene level AE data and ANEVA-DOT results in MDM cohort is given is supplementary **Data S1**.

## 1 Methods

### 1.1 Effect of a single regulatory variant on total gene dosage

Let us first consider a gene where its expression is affected only by one a single regulatory variant *v*. Following the notation in (*15*) let us assume that v has two alleles in the population: v_0_ and v_1_, and the associated allelic fold change (aFC) *δ*_1,0_, meaning that v_1_ induces *δ*_1,0_ times higher gene expression that the v_0_. The choice of allele labels is arbitrary and does not affect the downstream calculations. The expression of the gene, *e*_〈.〉,〈.〉_, in the three possible genotype groups are

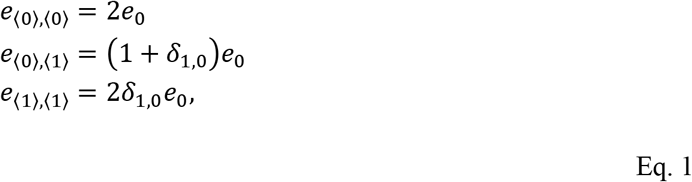

where *e*_0_ is the expression of the gene from a single haplotype carrying v_0_ (*15*). Assuming that *e*_〈0〉,〈0〉_ is the “normal” expression of the gene, departure of expression in the heterozygous individuals is

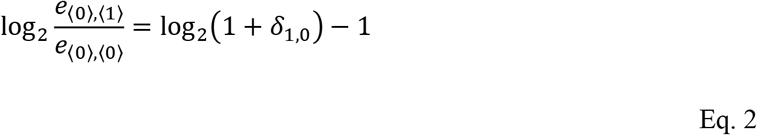

and in cases homozygous for allele v_1_ is

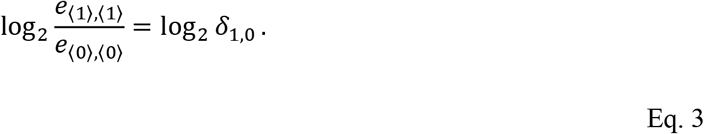

These functions are plotted for a range of aFCs in **Figure 1G**.

### 1.2 Analysis of Expression VAriance (ANEVA)

ANEVA couples a generative model for AE population data with the mechanistic link connecting the effect size of regulatory variants to variation in total gene expression, in order to derive estimates of genetic variation in gene expression (**Fig S1A**). In this section we describe different aspects of the models and concepts used for ANEVA.

#### 1.2.1 Expected genetic variation in gene expression (V^G^)

In this section we assume all the variants regulating a gene, their respective allelic fold changes, as well as their joint population frequencies as known. While these are typically unknown, this model is the theoretical basis that is helpful for understanding the model and for performing simulation studies. We aim to quantify natural variation in gene expression due to genetic factors in a population. First, we consider the case of a gene regulated by only one variant in the population presented in **Eq. 1–3**. Substituting the genotype-specific expressions, *e*_〈.〉,〈.〉_ from **Eq. 1**, and v_1_ allele frequency *p*, the expected deviation is

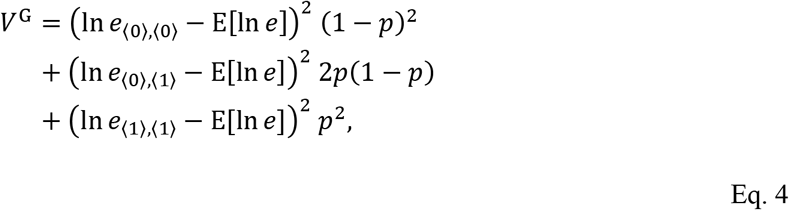

where the expectation E[ln *e*] is the expected log expression of the gene across all genotypes in the population:

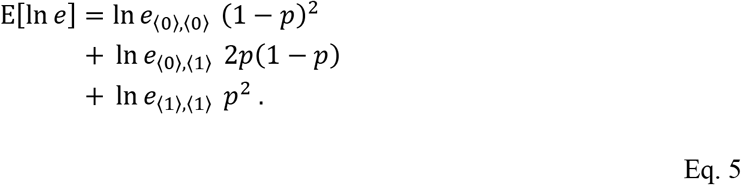

In general, the expected deviation can be calculated by averaging over all genotypes weighted by the respective frequencies in population:

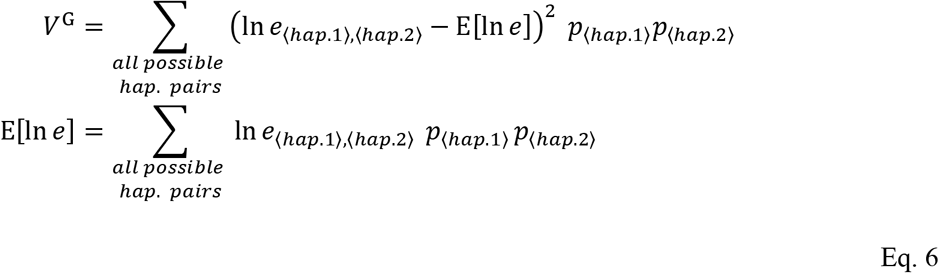

for the case where expression of a gene is regulated by *N* different variants v_1_, v_2_,… v_N_, with *m*_1_, *m*_2_,…, *m_N_* alleles in the population, respectively, by expanding *e*_〈hap.1〉〈hap.2〉_ in **Eq. 6** (see (*15*) for derivation), expected genetic deviation in expression is

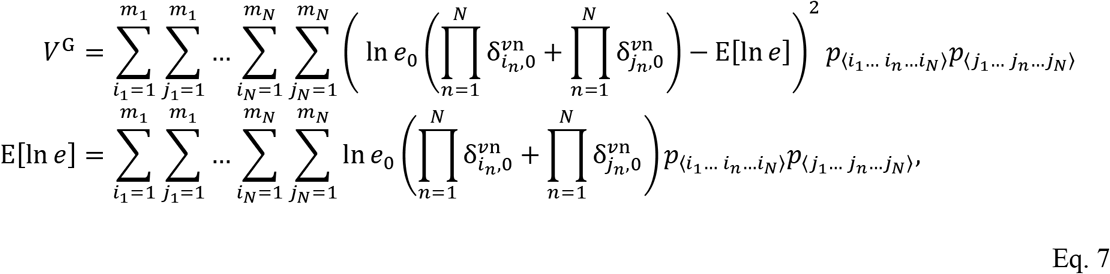

where 〈*i*_1_… *i_n_*… *i_N_*〉 denotes a haplotype carrying the *i*_n_-th allele of the *υ*n, and 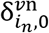 is the allelic fold change associated with allele *i*_n_, at the *n*^th^ eVariant *υ*n versus its reference allele 0, *e*_0_ is the reference expression associated with the case where the haplotype carries reference alleles for all eVariants (i.e. *e*_〈0… 0…0〉_). It is easy to show that *σ*^G^ does not depend on the reference expression *e*_0_. The haplotype probabilities, *p*_〈*i*_1_… *i_n_*…*i_N_*〉_ and *p*_〈*j*_1_…*j_n_*…*j_N_*〉_ depend on individual allele frequencies and the LD structure, and in the absence of LD will factorize into product of individual allele frequencies. The above equations can be used to simulate genetic variation in gene expression using an arbitrary set of regulatory variants. However, in practice, estimating genetic variation using this formulation is limited by the fact that for each gene usually only one or a few strong and common regulatory variants are available in known eQTL catalogs. More importantly, genes that are most intolerant to regulatory variation are expected to lack common regulatory variants identifiable as eQTLs, and for genes where regulatory variation is driven mainly by rare variants, eQTL-based estimates would be lacking or inaccurate.

#### 1.2.2 Distribution of AE data given complete genotype information

Effect of *cis*-regulatory variant on expression is reflected in both total expression and AE data (*15*). First we consider the case of a gene regulated by a single biallelic regulatory variant with a high expression allele, *H*, and a low expression allele, *L*. Let us assume that allelic expression is measured at a biallelic SNP (aeSNP) with reference allele, *R*, and alternative allele, *A*. AE data is only available in individuals that are heterozygous for the aeSNP. Thus, on average AE data at a given aeSNP is available in 2*p_R_*(1−*p_R_*) ≤ 50% of the population sample, where *p_R_* is the population frequency of allele *R*.

We denote the genotype of each haplotype as 〈*Regulatory allele; aeSNP allele*〉, and denote each individual using a pair of haplotypes. For example 〈*L; R*〉, 〈*H; A*〉 corresponds to an individual that is carrying the low expression regulatory allele, *L*, and the reference aeSNP allele, *R*, on one haplotype and the high expression regulatory allele, *H*, and the alternative aeSNP allele, *A*, on another (The order of the two haplotypes and also the choice of reference allele for the aeSNP is arbitrary and does not affect any of the analyses presented). As long as aeSNP alleles and the regulatory alleles are not in full LD, the allelic expression ratio across individuals in a population for such a gene forms a trimodal pattern consisting homozygous individuals (genotypes 〈*H; R*〉, 〈*H; A*〉 and 〈*L; R*〉, 〈*L; A*〉) with no allelic imbalance, and heterozygous individuals (genotypes 〈*H; R*〉, 〈*L; A*〉 and 〈*L; R*〉, 〈*H; A*〉) who show allelic imbalance skewed towards one or the other allele (**Figure 1A-D**). Specifically, true reference allele ratio, *r*, in each genotype group is:

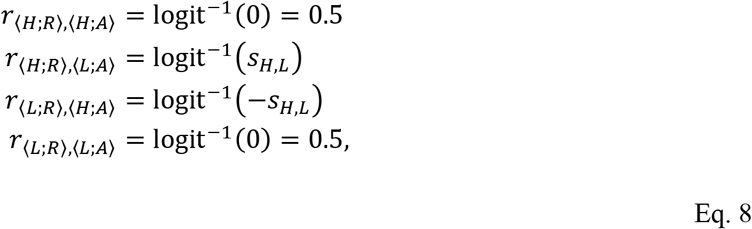

where *s_H,L_* is the effect size of the regulatory variant expressed as log aFC between the higher and the lower expressed alleles (*s_H,L_* = ln *δ_H,L_*; see **Eq. 1**), and the inverse-logit or the logistic function, logit^−1^, is defined as logit^−1^(*x*) = 1/(1 + *e^−x^*). Here we distinguish the *true reference allele ratio* from *observed reference allele ratio* that is obscured by binomial sampling noise in count-based AE data. The frequency of each genotype group in AE data is determined by the reference allele frequency of the regulatory variant as well as its linkage to the aeSNP. Specifically

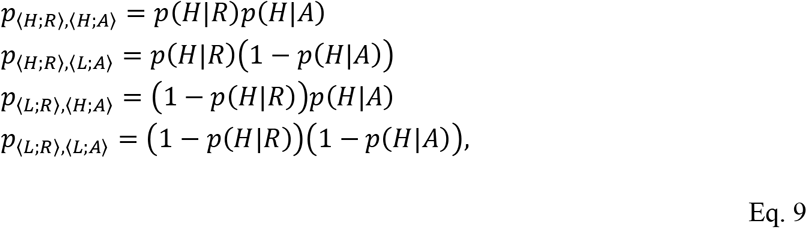

where *p*(*H*|*R*) and *p*(*H*|*A*) are the probabilities of having the high expressed allele of the regulatory variant on a haplotype carrying the reference or the alternative allele for aeSNP, respectively. In the absence of linkage disequilibrium between the regulatory and the aeSNP the two probabilities are equal to the reference allele frequency of the regulatory allele (*p*(*H*|*R*) = *p*(*H*|*A*) = *p*(*H*)).

In the general case where expression of a gene is regulated by *N* independent variants v_1_, v_2_,… v_N_, with *m*_1_, *m*_2_,…, *m_N_* alleles in the population, respectively, allelic expression will have 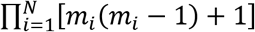 modes of imbalance in population (3^*N*^ if all regulatory variants are biallelic). The true reference allele ratio for each genotype group and its corresponding frequency in the population is

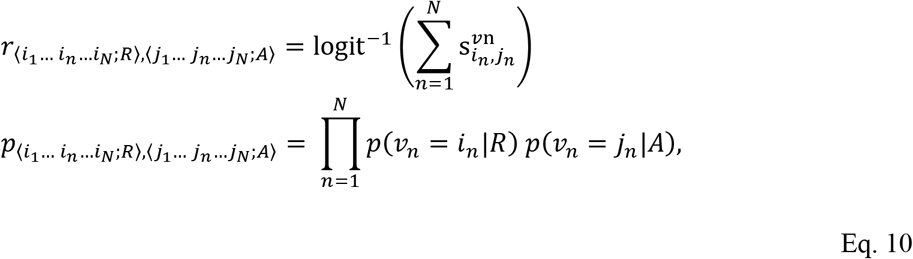

where 〈*i*_1_… *i_n_*… *i_N_*〉 and 〈*j*_1_… *j_n_*… *j_N_*〉 are the set of present alleles collocated on the same haplotype with *R*, and *A* alleles, respectively (*15*). The observed reference allele count in AE data, *y_t_*, given the sample genotype 〈*i*_1_… *i_n_*… *i_N_;R*〉, 〈*j*_1_… *j_n_*… *j_N_;A*〉 is a binomial sample drawn using the true reference allele ratio:

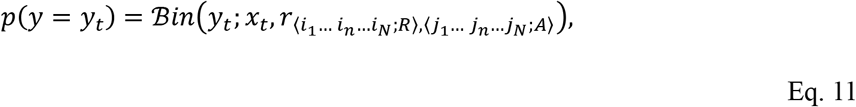

where *x_t_* is the total read count at the aeSNP. In practical cases, the full regulatory information is never available and the observed variation in reference allele counts exceeds that of a binomial distribution. In the next section we derive reference allele count distribution given limited regulatory variant information.

#### 1.2.3 Distribution of AE data given partial regulatory variant information

In the previous section, we introduced a model that describes patterns of reference allele ratio in a population under an ideal scenario in which every regulatory variant is known (**Eq. 10**). This theoretical model is too complex for practical applications where only a subset of regulatory variants affecting a gene are characterized. In this section, we derive a probabilistic model family for allelic expression population data under a realistic scenario in which only a subset of the regulatory variants is included in the model. Thereby, providing a consistent framework for modeling allelic expression data at various levels of complexity depending on the number of regulatory variants that are included explicitly in the model. This model is necessary in practical applications where limited data size does not allow for parameter inference beyond a certain level of model complexity, or when only a limited number of regulatory variants are characterized.

Let us assume a gene with *N cis*-regulatory variants that is described in **Eq. 10**. Here we assume only *M* variants are characterized, where 0≤*M*≤*N*. The true reference allele ratio for each genotype group and its corresponding frequency in the population is

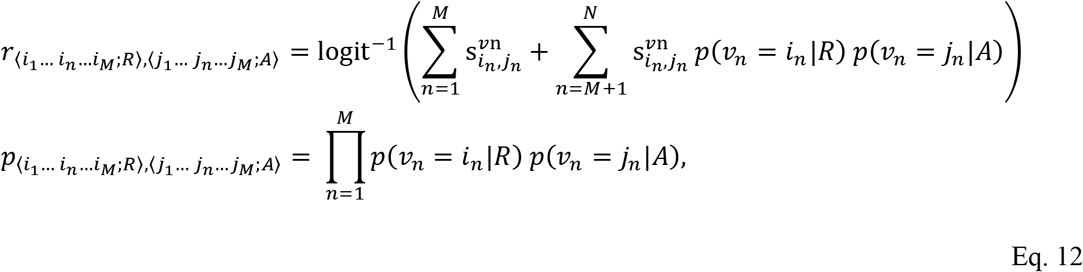

where only the first *M* variants are explicitly included in the model. The reference allele ratio in this case is comprised of two components: The total log aFC of the known variants 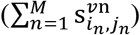, and the residual aFC, *s_r_*, that is the expected regulatory effect of the unknown variants 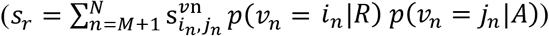. Assuming that all variants, *v*_M+1_ to *v*_N_, have finite effect sizes, according to the central limit theorem the expected effect of the unknown regulatory variants approaches normal distribution when the number of unknown variants is large (*N* − *M* → ∞):

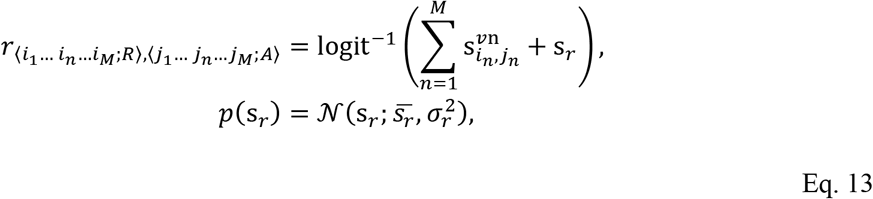

where 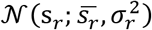 is the normal probability density function (PDF) evaluated at *s_r_* using unknown mean 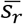, and variance 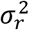 for the effects of the residual regulatory variants *v*_M+1_ to *v*_N_ in log aFC (In the absence of LD, 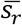 is zero). Using properties of normal PDF and applying logit transformation on both sides:

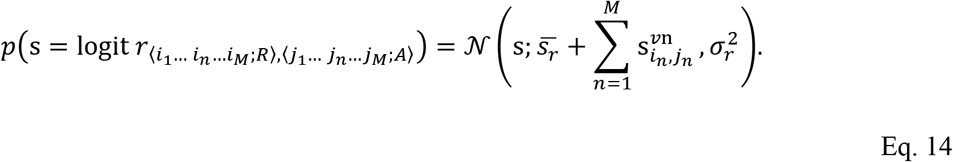

Therefore, the true reference allele expression ratio of a gene regulated by *N* regulatory variants given the partial genotype information from *M* known variants in a given individual is logit-normal distributed:

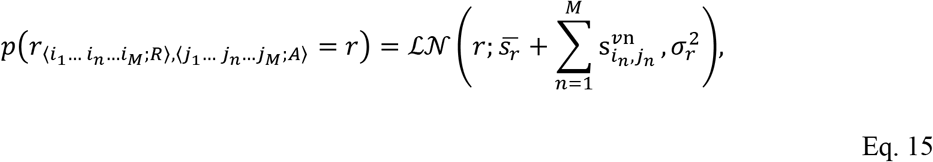

where, for any *r* between 0 and 1, the logit-normal PDF is defined as

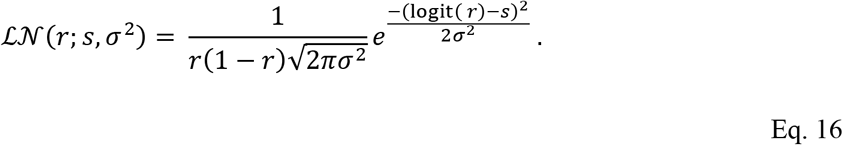

Recalling **Eq. 11**, the observed reference allele count in AE data is a binomial sample drawn using the true reference allele ratio. Since in this case *r*〈*i*_1_… *i_n_*…*i_M_;R*〉,〈*j*_1_… *j_n_*… *j_M_;A*〉 is given by a probability distribution:

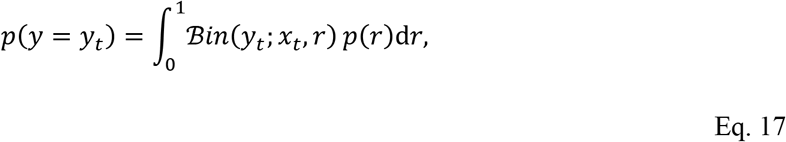

where *y_t_*, and *x_t_*, are the reference allele and the total counts at the aeSNP in a given sample *t*. Substituting *p*(*r*) from **Eq. 15**:

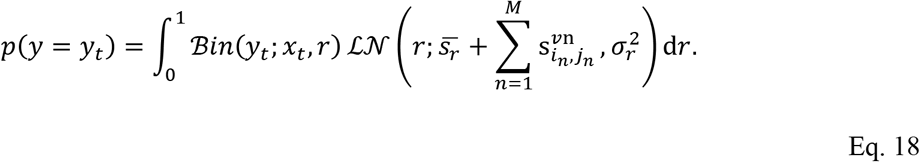

This integral is the Binomial-Logit-Normal (BLN) distribution:

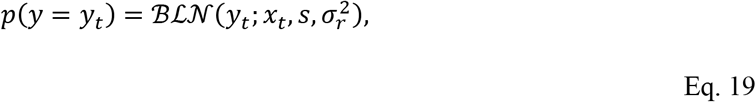

where 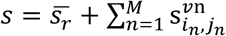 is the sum of regulatory difference between genotypes of the known 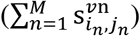, and the expectation for that of unknown variants 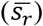 in log aFC, and 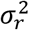 is its variance. We provide a general implementation of BLN probability density function, quantile function, and random number generation function in an R package available on GitHub: https://github.com/PejLab/bln.

#### 1.2.4 Population model for AE data underlying ANEVA

Current population AE data rarely shows more than three distinctive modes of allelic ratio. This is likely combined result of the properties of the regulatory landscape, noise and sample size. Following this intuition, we use the general model for true reference allele ratio in **Eq. 19** for *M* = 1 and a large, unknown, value for *N*. We further assume that there is no systematic regulatory discrepancy between the two haplotypes 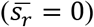. Thus, we model observed reference allele ratio of a given gene in a population by explicitly accounting for one latent biallelic regulatory variant with a distinctively large effect, and implicitly modeling the remaining variants using the distribution of their net regulatory effect. Accounting for genotype uncertainty associated with the latent variant, the final model for the reference allele counts is a constrained BLN-mixture model that is detailed in the following section.

##### Model description

Let us assume an aeSNP within a given gene. The aeSNP has two alleles in the population arbitrarily labeled as the reference (*R*), and the alternative allele (*A*). Allelic expression of the gene is measured in a set of individuals heterozygous for the aeSNP; multiple aeSNPs per gene are treated separately and summarized later. The input data to the model consists pairs of *x_t_*, and *y_t_* that are the total expression count and the reference allele expression count in the *t^th^* individual, respectively. Let us assume there exists a *cis*-regulatory variant, herein the *latent variant*, with a distinctive effect, strong enough to drive the general pattern of allelic ratios in the population data (Later on we relax this assumption and consider cases where there is no such variant). We refer to this variant as latent since its genotype is not explicitly present in our input dataset. The latent variant can be any *cis*-acting variant and is assumed to have two alleles in the population: the higher expressed allele (*H*), and the lower expressed allele (*L*). The reference allele expression count at the aeSNP, *y_t_*, is described by a constrained three-component BLN mixture likelihood model (**Fig S1B**) with parameters set *θ*:

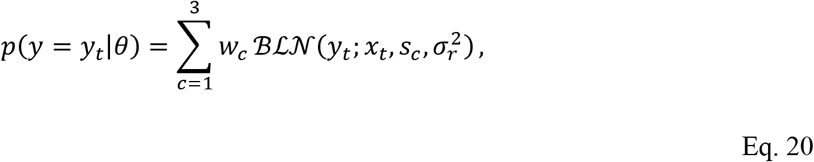

where the first component, *c* = 1, corresponds to individuals heterozygous for the latent variable, that carry the higher expressed allele, *H*, and the reference aeSNP allele, *R*, on the same haplotype (Genotype: 〈*H; R*〉, 〈*L; A*〉). The second component, *c* = 2, corresponds to individuals homozygous for the latent allele (Genotypes: 〈*H; R*〉, 〈*H; A*〉 and 〈*L; R*〉, 〈*L; A*〉), and the third component, *c* = 3, corresponds to individuals heterozygous for the latent variable, that carry the lower expressed allele, *L*, and the reference aeSNP allele, *R*, on the same haplotype (Genotype: 〈*L; R*〉, 〈*H; A*〉). Thus, allelic ratio *s_c_* in the three classes are

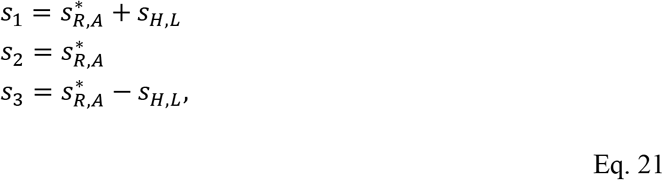

where *s_H,L_* is the log aFC associated with the latent variant (see **Eq. 8**), and 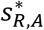 is log odds ratio of the a read aligning to the reference aeSNP allele to that of the alternative allele due to allelic mapping bias. From an analytical point of view, reference allele alignment bias acts as a spurious *cis*-regulatory effect. The log aFC associated with reference bias is given as 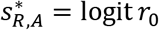 where *r*_0_ is the reference allele ratio in absence of *cis*-regulatory difference between the two haplotypes. Note that, 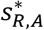 can be positive or negative, but *s_H,L_* is positive by definition as it describes the log ratio of the higher expressed allele to that of the lower expressed one. The component weight, *w_c_*, is proportional to the genotype frequencies (See **Eq. 9**):

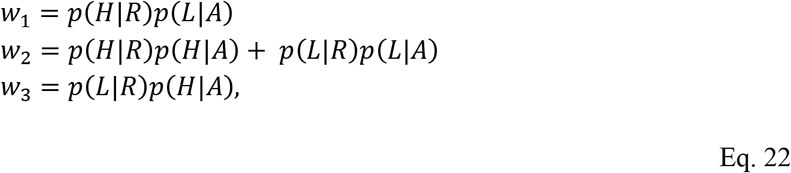

where *p*(*H*|*R*) and *p*(*H*|*A*) are the probabilities of having the *H* allele for the latent variant on a haplotype carrying the *R* and *A* allele for aeSNP, respectively. In the absence of LD these two conditional probabilities are equal. Additionally, *p*(*L*|*R*) = 1 − *p*(*H*|*R*), and *p*(*L*|*A*) = 1 − *p*(*H*|*A*) since the latent variant is assumed to be biallelic.

##### Reduced model variations and model selection

The full population AE model presented in **Eq. 20, Eq. 21**, and **Eq. 22** contains four free parameters: latent variant effect size (*s_H,L_*), its conditional allele frequencies (*p*(*H*|*R*) and *p*(*H*|*A*)), and the concentration parameter associated with residual regulatory effects (*κ*). We assume that the reference allele bias, 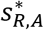, is given (see below how this parameter is set). In order to avoid over-fitting we considered three nested versions of the model with reduced complexity: 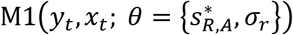 is the simplest of the three models that assumes there is no distinctive latent regulatory variant that dominates the population pattern in AE data. In this model *p*(*H*|*R*) = *p*(*H*|*A*) = 0 and consequently *s_H,L_* is excluded from the data likelihood. This model corresponds to cases where population AE pattern resembles a single cloud of beta-binomially distributed allelic counts centered around 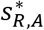 (*w*_1_ = *w*_3_ = 0; **Figure 1A-B**). 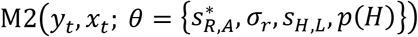 allows for a latent variant but assumes that there is no linkage disequilibrium between the latent variable and the aeSNP alleles (*p*(*H*|*R*) = *p*(*H*|*A*) = *p*(*H*)). This model corresponds to cases where population AE pattern forms three clouds of beta-binomially distributed allelic counts in each: one centered around 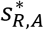 representing individuals homozygous for the latent variable, and two symmetrically placed on the two sides of the former, representing the heterozygous cases (*w*_1_ = *w*_3_). Finally, 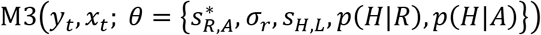 is the full model that allows for linkage disequilibrium between the two variants. This model corresponds to cases similar to those described by M2 but allowing for significantly different population frequencies in the two off-diagonal clouds (**Figure 1C**). In cases with full LD between the aeSNPs and the latent variant this model describes population patterns which includes only one cloud with average allelic imbalance significantly different from the 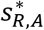 (**Figure 1D**). The model with optimal complexity was selected using Bayesian Information Criterion (BIC):

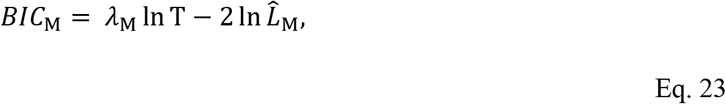

where *λ*_M_ is the number of free parameters learnt in model M ∈ {M1, M2, M3}, and 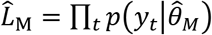 is the total data likelihood at the maximum likelihood estimates, 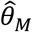, derived for model M and the likelihood model in **Eq. 20**. The model with the lowest BIC was selected at each aeSNP.

##### AE data preparation and parameter inference

The GTEx v7 data consists of 10,361 RNA-seq samples spanning 48 tissues and 620 individuals with whole genome sequencing data. The processing and quality control of these data is described in https://gtexportal.org, and the allelic expression data was produced as described in (*17*). Briefly RNA-seq reads were aligned using STAR in two-pass mode, and allelic counts were generated from uniquely mapping reads using the GATK ASEReadCounter tool with the following settings: *-U ALLOW_N_CIGAR_READS -minDepth 1 --allow_potentially_misencoded_quality_scores --minMappingQuality 255 --minBaseQuality 10*. aeSNPs in low mappability regions, those with potential genotyping error, and those having large allelic mapping bias in simulations were discarded as proposed in (*24*). We used all aeSNPs that are expressed at least 30 reads in at least 6 individuals in a given tissue in GTEx v7 data. This yielded median of 43,160 aeSNPs per tissue. Reference allele bias, 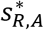, at each aeSNPs was fixed to the mean reference allele ratio in each sample for each of the 16 possible combinations of reference and alternative allele as in (*16*) (*24*). For all three models, M1-M3, we used gradient descent (Matlab function *fmincon*) to minimize negative log-likelihood function of **Eq. 20** over all the individuals with data available at a given aeSNP:

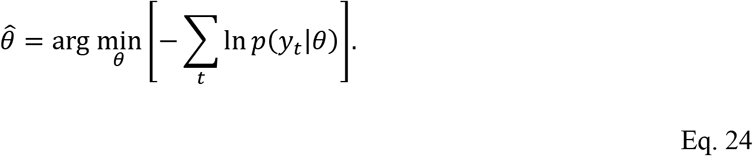

Standard deviation of the residual effects, *σ_r_*, was optimized within [10^−4^, 1.5]. The 99% probability mass of 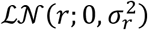 allows for 1.0005-, and 1074-fold regulatory difference between the residual effects of two haplotypes at the lower (*σ_r_* = 10^−4^), and the upper bound (*σ_r_* = 1.5), respectively. Latent variant log aFC, *s_H,L_*, was optimized within [0, ln 100] range.

For fitting ANEVA model on GEUVADIS consortium data, we used the same preprocessing and parameters setting as for the GTEx data. We used AE counts from 421 individuals including CEU (*n* = 77), FIN (*n* = 89), GBR (*n* = 86), TSI (*n* = 92) and YRI (*n* = 77) subpopulations. ANEVA was once calculated on the entire dataset, and once using each subpopulation independently.

##### Model identifiability in the absence of LD

In the no LD model, M2, allele frequencies for the latent variant alleles are only identifiable up to their minor allele frequency. By definition the two heterozygous components in the model are symmetric with regard to their allelic imbalance 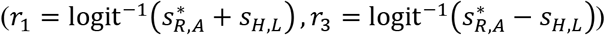. The problem arises in the absence of LD, where the two components also have the same expected population frequency (*w*_1_ = *w*_3_) and the likelihood function is symmetric around 50% allele frequency for the latent variant. In these cases for any MLE parameter sets 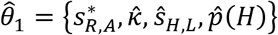 there is another set 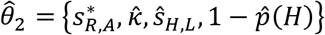 that has an equal data likelihood:

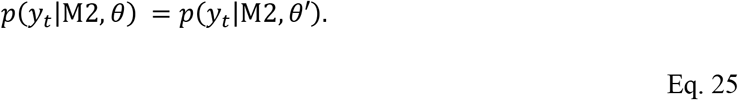

Thus, learning the model parameters for M2 requires an additional step to narrow the maximum likelihood solution down to either 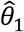 or 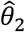, in other words, to find out whether the higher expressed allele is the major or the minor allele.

We used a Gaussian mixture model on the total expression at the aeSNP to compare evidence for 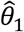 and 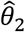. This model was constrained to incorporate inferred values from the AE data including the effect size and the minor allele frequency of the latent variant. Additionally, the total expression model is further constrained by partial information about the genotype of each individual. Specifically, individuals belonging to the second component in the AE model (*c* = 2; See **Eq. 20**) are assumed to be homozygous, and the rest to be heterozygous, for the latent variant. Let us assume *p_H_* to be the frequency of the higher expressed allele in the population. The frequency of individuals homozygous for the latent variant within all homozygous individuals is

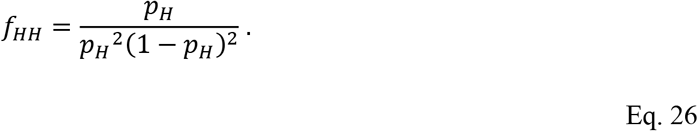

Given the expected total gene expression of a gene regulated by a single regulatory variant presented in **Eq. 1**, the total expression model, F(*z_t_, c_t_*; *ψ* = {*s_H,L_, p_H_, e_L_, σ*^2^]) was defined as:

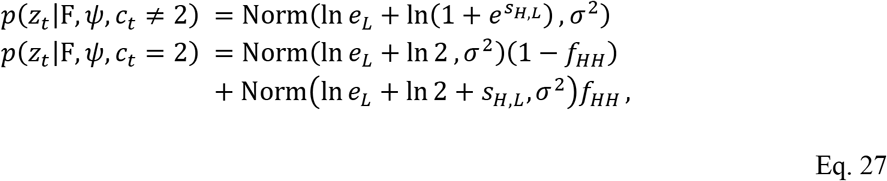

where *z_t_* is the library normalized expression in the *t*^th^ individual in the logarithmic scale, *c_t_* is the maximum a posteriori class associated with the *t*^th^ individual in the AE population model (**Eq. 20**). All model parameters are fixed except for *e_L_*, the expression for the lower expressed allele, and the noise variance *σ*^2^. We used an EM-like algorithm to recursively maximize the data likelihood over all positive values for *e_L_* and *σ*^2^ under two scenarios: first, assuming 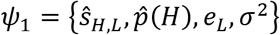, and next, assuming 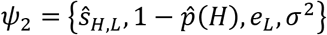, where *ŝ_H,L_* and 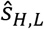 are the MLE estimates from the AE model (**Eq. 24**):

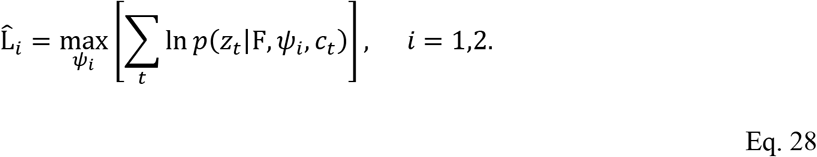

For each pair of MLE solutions 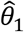 and 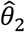 for the AE model M2, we then selected the one that corresponds to higher data likelihood, 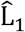 or 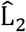, in the total expression model F (**Eq. 27**).

#### 1.2.5 Estimating V^G^ from AE population model parameters

The AE population model corresponds to a gene that is regulated by two regulatory variants: one biallelic variant that is the main variant with alleles H and L, and one infinite allelic variant that captures the residual regulatory variation *s_r_*. Thus, to calculate V^G^ we used the MLE estimates *σ_r_, s_H,L_, p*(*H*|*R*), *p*(*H*|*A*) from the AE model (**Eq. 24**) and aeSNP allele frequencies *p*(*R*) and *p*(*A*) in **Eq. 7** and **Eq. 10**:

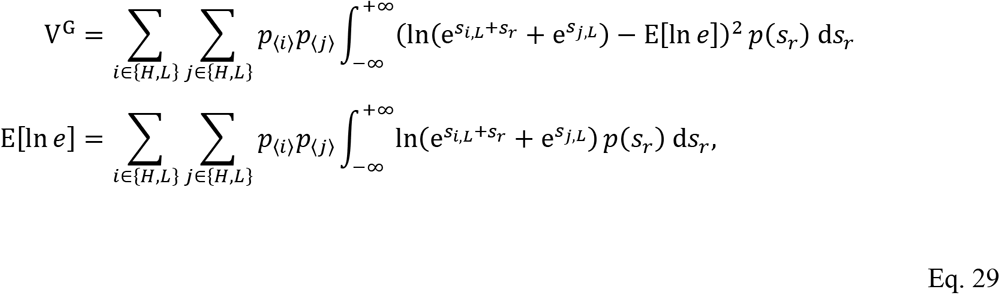

where

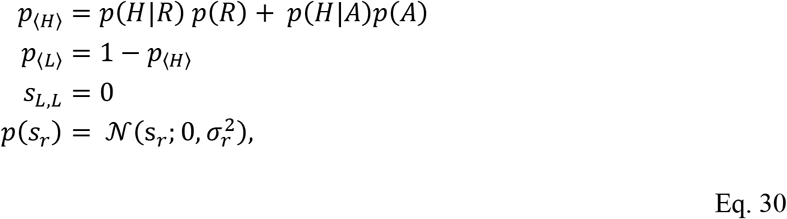

and reference expression *e*_0_ is set to 1 as its value does not affect V^G^.

#### 1.2.6 Combining V^G^ estimates across aeSNPs and tissues

ANEVA model was fit on SNP-level data. In order to derive gene-wise V^G^ estimates we had to combine several SNP level results. Similarly, we had to combine V^G^ estimates from different tissues to derive a single cross-tissue 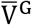 estimate that can be compared to genome-derived external scores which are not tissue-specific. To do these, we used weighted harmonic mean. The choice to use harmonic mean instead of arithmetic mean was motivated by the fact that we are inherently interested in genetic precision, the reciprocal of variance, of regulation. In other words, in this analysis we are most interested in cases where V^G^ is the smallest. Additionally, due to the nature of the AE data and the model large V^G^ estimates tend to be noisier than the smaller ones which require high counts and large number of samples. Harmonic mean allows for higher emphasis on smaller V^G^ values. Additionally, V^G^ estimates from lower count SNPs, or tissues with lower expression for a given gene are likely to be less biologically relevant. To take this into account the total number reads supporting an SNP-level estimate, and the tissue-wise expression of a gene were used as the weight for each estimate in harmonic mean used for collapsing SNP-level and tissue-specific estimates respectively.

#### 1.2.7 Simulating the effect of read coverage in V^G^ estimates

To evaluate the effect of read coverage in V^G^ estimate we used a down-sampling experiment. To do this we generated a total of 768 down-sampled sets of AE data. We selected eight genes to span over a range of SD^G^ values. To allow for thorough down sampling we used genes with a high original read coverage (average coverage over 500 reads across all GTEx individuals). For each gene we down-sampled the total read count at 100%, 95%, 90%,…, 5% and at 4%, 3%, 2%, and 1% of the original total coverage. At each coverage level we produced down-sampled AE data using binomial sampling from the reduced coverage and the original reference allele ratio. We repeated this procedure four times to have four down-sampled datasets from each gene and each reduced coverage level. We then ran ANEVA on these data and calculated V^G^ (**Fig S4A**). As expected, the reduction in the converge progressively increased noise in the V^G^ estimates in all tested genes. However, we did not observe any consistent pattern of correlation between V^G^ estimates and the coverage in this simulated experiment, showing that while V^G^ estimates from low read counts may be noisier, they are not systematically biased towards high or low values.

#### 1.2.8 Comparing V^G^ estimates to eQTL data and local heritability of expression

To compare AE based estimates of V^G^ to eQTL-based estimates we used aFC effect size estimates from the most significant eQTL for each gene in each tissue (*15, 17*) available for download at: https://gtexportal.org/home/datasets). When available the effect size and the allele frequencies for the eQTL was used in **Eq. 4** to calculate V^G^.

To estimate heritability using ANEVA we used the ratio between V^G^ and the total expression variance, V^T^. To calculate V^T^ we aimed to use the most straightforward approach to avoid any spurious correlation with external heritability estimates caused by dependencies in V^T^. For each tissue, we used median expression of genes across all individuals as the reference and used all genes with median expression over 250 reads to normalized samples for median expression fold change vs. the reference as described before (*25*). BSLMM estimates of heritability were derived from GTEx whole blood samples, and well as DGN data (*26*) as described by Wheeler et al. (*18*). For calculating the correlation between different methods, we used genes that were shared by all (*n* = 1939).

#### 1.2.9 Other datasets and gene lists used for comparison to V^G^

We obtained pLI (*1*), RVIS (*5*), ncRVIS and ncGERP (*27*) estimates of constraint from the respective papers without any additional filtering or transformations. We defined enhancer size for each gene based on the GeneHancer predictions of enhancers and their target genes (*21*). We downloaded the file https://genecardsdata.blob.core.windows.net/public/GeneHancer_version_4_4.xlsx and further filtered the data to include only enhancers with an enhancer score ≥0.7 and enhancer-gene link score ≥7. Some of the scores in **Figure 3C** were multiplied by −1 to visualization purpose. This is reflected in corresponding labels. Spearman correlation was reported, and 95% confidence intervals were calculated using the bias-corrected and accelerated bootstrapping.

The genesets used for **Figure 3D** are provided in **Table S3**. Specifically, Autosomal dominant, Autosomal recessive set was taken from OMIM (https://www.omim.org, downloaded in Feb. 2019). Haploinsufficient gene list includes genes marked as haploinsufficient with score 2 or 3 in ClinGen Dosage Sensitivity Map (https://www.ncbi.nlm.nih.gov/projects/dbvar/clingen/, downloaded in Sept. 2018). LoF tolerant and LoF intolerant genes were defined as genes with pLI ≤ 0.1, and pLI ≥ 0.9, respectively (*1*). Nonessential, Dosage sensitive gene lists were taken from the copy number variation map of the human genome by Zarrei et al. (*28*). Mutated in autism gene list were genes with gene-score 1 or 2 in SFARI human gene module (https://gene.sfari.org/database/human-gene/SFARI, Nov. 2018 release). Mutated in CHD set is taken from (*29*). Mutated in DD set is taken from (*30*). Mutated in MDM set is taken from (*23*). The remaining sets associated with common phenotypes and diseases were all taken from GWAS catalog (https://www.ebi.ac.uk/gwas/docs/file-downloads, Nov. 2018 release).

#### 1.2.10 Implementation and availability of ANEVA

ANEVA is implemented in a MATLAB package available on Github: https://github.com/PejLab/ANEVA. As input, it takes variant level AE data for a set of individuals in a population, as well as the allele frequency for aeSNPs. The package uses these data to fit the population model at each available aeSNP, calculated the expected V^G^ and combine them over genes. We provide sample data for input files. This package can be used for calculating V^G^ estimates from new reference datasets. Additionally, we provide pre-calculated V^G^ estimates for current and future datasets available to us here: https://github.com/PejLab/Datasets/tree/master/Reference_Vg_Estimates

### 1.3 ANEVA Dosage Outlier Test (ANEVA-DOT)

ANEVA-DOT is a method that takes one individual at a time and inspects allelic expression for each of the genes of the individual in the context of the general population variation V^G^ for that gene. Let *y_g_* and *x_g_* be the reference allele count and the total count at a given aeSNP representative of allelic expression of gene g with natural genetic variation, 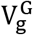, in its dosage. Under the null hypothesis for ANEVA-DOT, observed reference allele count *y_g_* is described by total coverage (*x_g_*), natural genetic variance in dosage 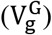, as well as other practical considerations, namely, reference bias and substitutional noise. First, we describe key idea behind ANEVA-DOT using a simplified version that only includes *x_g_* and 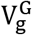. Next, we provide the full test that takes the two additional practical issues into account.

#### 1.3.1 Simplified ANEVA-DOT

Let us assume a regulatory mutation M that changes the overall gene dosage by *Δ*_M_ = ln *e*_〈0〉,〈M〉_ − ln *e*_〈0〉,〈0〉_ relative to the unmutated dosage *e*_〈0〉,〈0〉_. By definition 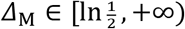 since a single *cis*-acting mutation can only disrupt one copy of the gene. While we refer to mutations for better intuition, the assumptions are appropriate for rare variants beyond de novo mutations. From **Eq. 2**

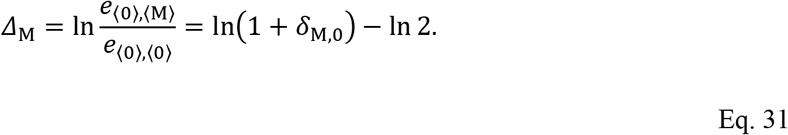

Solving this equation for *δ*_M,0_, the aFC associated with the mutation is

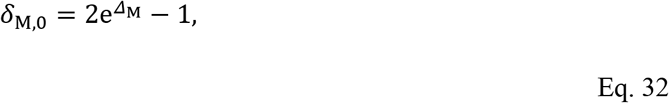

and the true allelic ratio of the mutant haplotype is

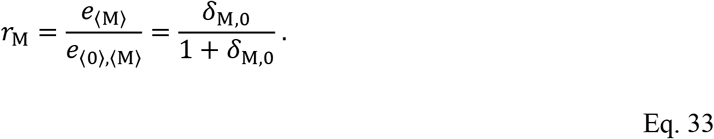

Therefore, probability of observing *y* reads from the mutant allele conditioned over *Δ*_M_ is

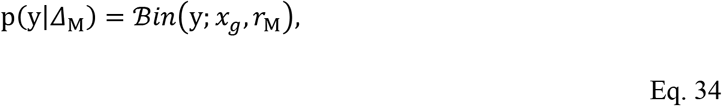

where *r*_M_ is given as a function of *Δ*_M_ as given in **Eq. 33**. Under the null hypothesis, we assume *Δ*_M_ is normal distributed with zero mean and variance 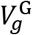 that is estimated by ANEVA from a reference population. Additionally, we bound *Δ*_M_ to fall within 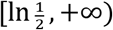:

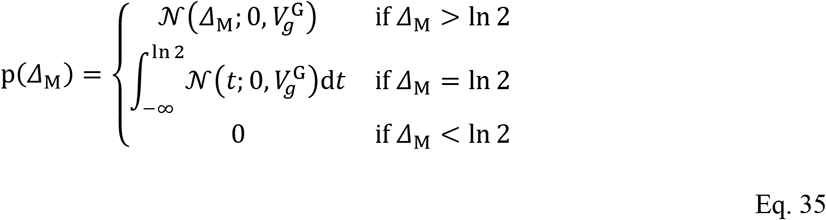

Thus, probability of observing *y* reads from the mutant allele is

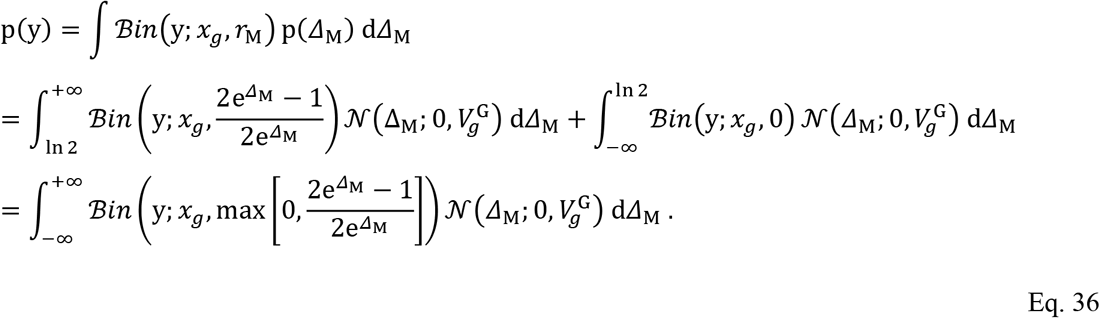

We sum the value on the two tails of the distribution for the mutant allele count *y* in **Eq. 36** and use the lighter tail to derive the test p-value as

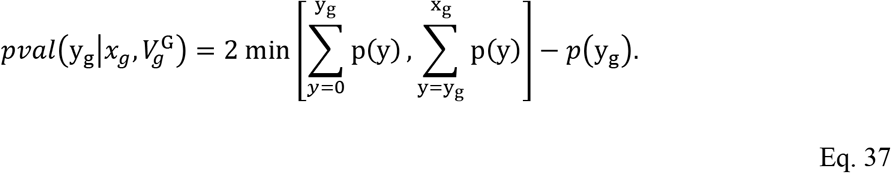

#### 1.3.2 ANEVA-DOT

Here we add two additional factors into the test: 1) substitutional noise probability (*π*), and 2) reference allele alignment bias. Specifically, *π* is the probability of a reference (or alternative) allele in the original RNA sample to appear as the alternative (or reference) in the data due to sequencing error or another similar source of noise along the process. The prevalence of this rare type of noise can be approximated in allelic expression libraries using the observed bases that are in conflict with the sample genotype (*18, 24*). Let us assume *r* to be the true reference allele ratio in a sample. Under this substitution model we expect *rπ* fraction of the reads to appear as the alternative allele due to noise. Conversely, we would expect (1 − *r*)*π* of the alternative allele expression to appear as reference. By including this the expected mutant allele expression in **Eq. 33** is adjusted as

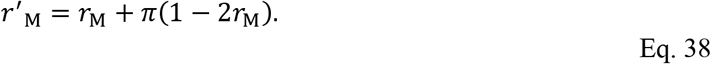

Next, we consider reference allele alignment bias that is when the allelic expression ratio is systematically skewed towards one of the alleles due to sequence alignment other artifacts. Several approached have been developed to alleviate this problem, and novel alignment approaches may resolve it completely (*31, 32*). However, since it is still present in most current data sets, it needs to be addressed. Let us assume a null reference allele ratio *r_0_*. We assume this bias in allelic expression is due to a multiplicative effect that does not depend on the expression itself. Log aFC associated with the reference bias is logit(*r*_0_). Using additivity of log aFC, the reference ratio after accounting for bias and noise is

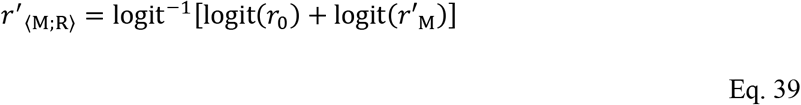

when the mutation *M* is on the same haplotype with reference aeSNP allele *R*, and

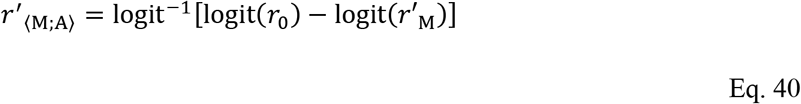

when it is on the haplotype carrying alternative aeSNP allele *A*. Assuming that the mutation is equally likely to occur on either haplotype, probability distribution in **Eq. 34** becomes the following mixture model:

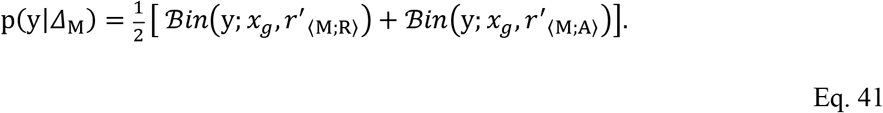

Thus, the probability of observing *y* reads from the mutant allele from **Eq. 36** changes to

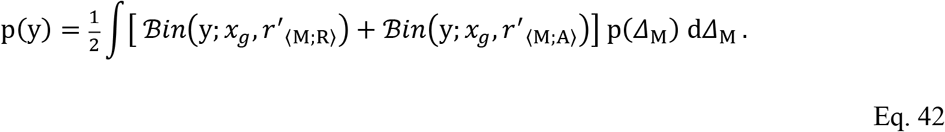

Probabilities calculated from this function is used in **Eq. 37** to derive ANEVA-DOT p-values.

#### 1.3.3 The assumptions and applications of ANEVA-DOT

ANEVA-DOT aims to find rare variants with unusually large regulatory effect. Based on this we assume that such a variant is, first, in heterozygote state, and second, far larger in effect size relative to any common regulatory variants for a given gene. The first assumption means the effect of the variant can be observed in AE data, and the second assumption allows us to calculate the expected effect on the total expression and compare it V^G^. These mean that ANEVA-DOT is not expected to capture cases where the combined effect of two high (or low) expressing regulatory variants on two haplotypes, either as homozygous variants, or compound heterozygotes, lead to an extreme gene dosage. Conversely, two rare haplotypes with multiple low and high expressing common alleles, respectively, could be paired and produce extreme AE. This pattern of common variation in principle can appear as an ANEVA-DOT outlier gene and provide a false positive hit of a dosage outlier. However, these situations will be very rare when V^G^ estimates are based on large population samples that represent all haplotypes of reasonable frequency in the population. Furthermore, in principle the BLN-based model family developed here for ANEVA, and the ANEVA-DOT model both allow one to directly include regulatory effect associated with all known regulatory variants in the individual of interest. However, implementing this model to integrate all available eQTL and AE data at the population and individual level involves overcoming several practical challenges that is beyond the scope of current work and it will be a goal for future research.

#### 1.3.4 Implementation and availability of ANEVA-DOT

ANEVA-DOT is implemented in an R package available on Github: https://github.com/PejLab/ANEVA-DOT. As input, it takes pre-calculated V^G^ reference estimates (provided for 48 GTEx tissues, and for GEUVADIS LCLs in **Table S1**), and the tested individual’s AE data either based on aeSNPs or summarized to haplotypes (*31*), in which case the ANEVA-DOT can be run without access to patient genotype data. Optionally, the input can include a vector of null allelic ratios, 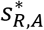, to account for the reference bias in each pair of allelic counts. If missing one library-wide estimate of 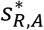 will be calculated using the median ratio between the reference and alternative allele counts using 20% of the data with the highest read coverage. We provide lower bounds for occurrence rate of significant ANEVA-DOT genes in all tissues available in GTEx v7 data in **Table S3**. These can be used to build blacklists for ANEVA-DOT analysis in difference tissues with appropriate thresholds to exclude genes prone to false positives. Additionally, we provide full summary statistics for ANEVA-DOT runs on the current and future reference datasets available to us in https://github.com/PejLab/Datasets/tree/master/ANEVA_DOT_frequencies.

### 1.4 ANEVA-DOT application to GTEx, GEUVADIS, and MDM disease samples

#### 1.4.1 GTEx

In order to first test ANEVA-DOT in the general population, we applied it to 515 skeletal muscle samples in GTEx v7 data preprocessed as described in section 1.2.4.3 *AE data preparation and parameter inference*. Fourteen individuals with an unusually low number of tested genes (*n* < 2888; threshold was defined as Q1 - 1.5IQR, where Q1 is the first quartile of the data and interquartile range, IQR, is the midspread defined as Q3-Q1) were removed from downstream analysis, in addition to 35 individuals with an unusually high rate of outlier genes, (*n* > 26 that is 0.79% outlier rate; threshold was defined as Q3 + 1.5IQR). Next, we discarded 113 genes which appeared as dosage outlier in over 1% of the individuals as prone to false positive (**Table S4**). To decide if a gene is a dosage outlier in over 1% of the individuals, we used the Clopper-Pearson method to calculate confidence intervals for the MLE estimates of the binomial distribution parameter, and discarded genes where the lower bound exceeded 1% rate (**Table S4**). After this filtering, each gene was on average an outlier in 0.4 individual in GTEx, and in turn, each individual had a median of 4.5 ANEVA-DOT genes. These filtering steps are illustrated in **Fig S8**.

To test for enrichment of putatively disruptive variants, rare variants were defined having a MAF less than 1%, both within GTEx and gnomAD release2.0.2. To test for enrichment of rare SNVs and indels near ANEVA-DOT outliers, we selected genes which were significant in at least one individual. Within this set of genes, we considered all rare variants within 10kb upstream of the TSS and within the gene body, in all available individuals. Individual-gene pairs that did not have a DOT score available were excluded from downstream analysis. All rare variants, across outlier and corresponding non-outlier individual-gene pairs, were annotated by their VEP consequence. In cases where an individual-gene pair had multiple proximal rare variants, each unique variant consequence was considered. Genes having both SNVs and indels with the same consequence were also considered in aggregate. For each group of rare variants considered, relative risk was defined as the odds ratio of an outlier individual carrying such a variant to that of a non-outlier individual:

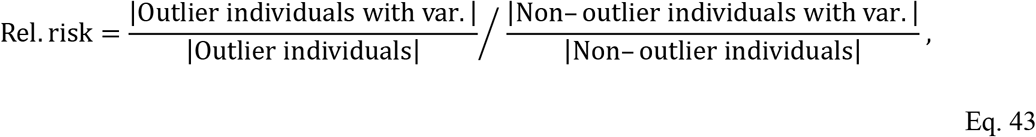

where |…| is the cardinality of the set. Confidence intervals for these relative risk estimates were reported, and significance was declared if the 95% confidence interval did not contain 1. Under the null hypothesis, a rare variant of a particular class does not influence a gene’s probability of emerging as an ANEVA-DOT outlier. We ensured that these results are replicated in unseen samples using other data that were not used in any of the prior analyses to estimate V^G^ (data not shown).

#### 1.4.2 GEUVADIS

For ANEVA-DOT analysis across subpopulation we used AE counts from 344 individuals including FIN (*n* = 89), GBR (*n* = 86), TSI (*n* = 92) and YRI (*n* = 77) subpopulations in GEUVADIS consortium data. We excluded CEU (*n* = 77) subpopulation from this analysis due to previously reported clonality issue in the cell lines (*33*).The data was preprocessed as described in section 1.2.4.3 AE *data preparation and parameter inference* similar to GTEx data. For comparison of the population effects presented in **Fig S10** we only considered AE data from aeSNPs with reliable genotype (genotype warning flag = false; (*24*).

#### 1.4.3 MDM cohort

The majority of the cohort of Mendelian neuromuscular disease patients was previously described in Cummings *et al*. Briefly, patients with available muscle biopsies were referred from clinicians from March 2013 to January 2019. The cohort includes samples previously diagnosed by WES/WGS (that were used as positive controls in Cummings *et al*.), samples that were diagnosed via RNA-seq, and the remaining genetically undiagnosed samples. The cohort described in this study includes 10 previously unpublished samples that were sequenced after the publication of Cummings *et al*. cohort. In total we used 70 samples. All samples are individually plotted in **Fig S11–S13**, and **Fig S15–S17**, and detailed in **Table S5**. Complete AE data and ANEVA-DOT results for MDM cohort are provided in supplemental **Data S1**.

AE data on all samples were generated as described in Cummings et al. using GATK ASE Read Counter package version 3.6 (*24*). Heterozygous variants that passed VQSR filtering were first extracted for each sample from exome sequencing VCFs using GATK SelectVariants. Sample-specific VCFs and RNA-seq BAMs were used as input to GATK ASEReadCounter, requiring coverage in the RNA-seq data of each variant to be at least 20 reads, with a minimum base quality of 10 and counting only uniquely mapped reads. The resulting AE file was annotated with the Variant Effect Predictor (VEP; (*34*)) version 85 against Gencode v19 using the Hail platform (https://hail.is/).

For samples where the genetic diagnosis was known, we first manually evaluated whether the gene affected by the disease-causing variant would be expected to exhibit allele imbalance (e.g. at least 1 loss-of-function variant) using the IGV browser (*35*). We then evaluated whether the gene harboring the pathogenic variant(s) were identified with ANEVA-DOT. For undiagnosed samples, the available DNA data was evaluated in the patient to identify putatively pathogenic variants using inheritance patterns (where parental data was available), functional annotation, and variant frequency in reference databases such as gnomAD (*2*). For each gene with a plausible rare variant, the splicing patterns in the available RNA-seq data were also evaluated. To evaluate the number of ANEVA-DOT significant genes in known MDM genes, we used union of a previously published list from: https://github.com/berylc/MendelianRNA-seq/blob/master/data/NMD_genes.list and all causal genes in the previously diagnosed samples to ensure this list includes all true positive genes. The final list that we used in our analyses is given in **Table S5**.

We used GTEx muscle tissue estimates of V^G^ to run ANEVA-DOT on the MDM cohort, and removed 113 false positive prone genes from the analysis as described for in section 1.4.1 *GTEx*. Out of our 70 samples in our cohort, five were excluded: two due to low quality RNA-seq data, and three due to presence of too many outlier genes by ANEVA-DOT which can occur due to sample contamination or genotyping issues (**Fig S17**). Of the remaining 65 patients, 32 had a genetic diagnosis from a combination of WES/WGS/RNA-seq, 11 for which we would not expect to exhibit allele imbalance due to the nature of the pathogenic variants (e.g. hemizygous loss of function variants on chrX, or in-frame pseudoexon insertions; **Fig S13**). Of the remaining 21 samples that were expected to exhibit allelic imbalance, in three cases the disease gene lacked AE data due to low expression or lack of aeSNP (**Fig S12A**), and in two cases the disease gene did not show any allelic imbalance (**Fig S12B-C**). Based on our further manual evaluation, one possible justification might be that in both cases, two loss-of-function mutation in *trans* may balance the NMD effect. In the remaining 16 samples, ANEVA-DOT identified the gene harboring the pathogenic mutations in all cases.

For benchmarking ANEVA-DOT we use binomial and beta-binomial tests of allelic imbalance. Binomial test was performed as a two-tailed test against the null hypothesis that AE data is binomial distributed with null reference ratio *r*_0_. The null reference ratio used was identical to what was used for ANEVA-DOT. Beta-binomial test was also a two-tailed test against the maximum likelihood beta-binomial fit for each individual’s AE data. All tests were corrected for false discovery rate using Benjamini-Hochberg procedure. Resulting genes with an FDR < 5% were further followed up in patient samples using the seqr variant interpretation platform (https://seqr.broadinstitute.org). All significant cases were evaluated regardless of prior disease association.

## 2 Supplementary texts

### 2.1 Supplementary text 1: Comparing AE and eQTL based estimates

We derived V^G^ estimates from lead eQTLs to ensure that our AE based estimates are concordant with what is expected from the much more standard eQTL data. The two sets of estimates were generally well correlated with median rank correlation 0.75 across GTEx tissues. For genes shared between the two methods, AE based estimates were slightly larger (median ratio 1.17), and comparable and slightly more correlated with other metrics (e.g. Spearman correlation with pLI of 31.7% vs. 29.3% and Spearman correlation with ncGERP 23.7% vs. 18.2%), and more consistent with our background knowledge (e.g. area under the ROC curve for classifying haploinsufficient genes of 65% vs. 56%, and for genes mutated in autism 67% vs. 61%). This was expected as the AE model is supposed to capture all regulatory effects on a gene versus the eQTL based estimates.

However, the eQTL-based estimates showed surprisingly high correlation to the local h^2^ estimates derived with BSLMM on an independent dataset, the DGN data (Spearman correlation of 59% vs. 32% for AE-based estimates; **Figure 2D**). Considering that the eQTL-based estimates of V^G^ do not appear to be particularly more consistent with any other external datasets that we used, and the fact that BSLMM estimates of local h^2^ on GTEx have low correlation to BSLMM estimates on DGN (Spearman correlation of 20%), we believe that this exceptionally high concordance is due to sparse linear regression model underlying BSLMM capturing mostly common regulatory variation (i.e. the eQTLs) in the current sample sizes. Thus, we expect this correlation to local h^2^ estimates from methods like BSLMM to decrease in future when they are applied to much larger transcriptome profiling datasets.

Another interesting aspect of comparing V^G^ estimates derived from AE and eQTL data is the subset of genes each of them cover. For instance, analysis of adipose subcutaneous GTEx tissue presented in **Fig S2** yields 7556 genes with V^G^ estimate when AE data is used and 7467 when eQTL data is used. However, only 3237 (~43%) genes have V^G^ estimates in both. Furthermore, the genes specific the eQTL-based analysis tend to have drastically larger V^G^ estimates. This is in clear contrast to the trend in the shared genes which AE-based V^G^ estimates are systematically larger (**Fig S2D-G**). This points to the fact that the genes covered by the two approaches are very different, which is due to a fundamental difference between ANEVA and eQTL analysis. ANEVA aims to quantify the genetic regulatory variation affecting a gene regardless of variant frequencies and effect sizes. This only depends on the availability of aeSNPs for each gene. On the other hand, eQTL analysis aims to identify when there is a significant regulatory variation signal present for a gene. Thus, by definition genes less tolerant to *cis*-regulatory variation tend to be absent in eQTL data. Since in our analysis we are most interested in genes intolerant of regulatory variation, we decided to use the AE-based estimates for all subsequent analyses. Additionally, since ANEVA-DOT is also done on AE data, learning ANEVA from AE data allows for our pipeline to account for some of the other biases present in AE data that we may not fully account for or exclude (e.g. those in low mappability regions) from our analysis. Combining AE and eQTL data for learning V^G^ estimates is not trivial but it is an intriguing possibility for future work as it would potentially allow for higher quality estimates that also span more genes.

### 2.2 Supplementary text 2: PPV and FPR derivation for ANEVA-DOT in MDM cohort

We found a total of 349 outlier genes by ANEVA-DOT in our 33 undiagnosed cases that passed quality control. Importantly, 12 of these patients had one or more known MDM outlier genes (total of 17 genes). We wanted to know what fraction of these genes identified by ANEVA-DOT in the undiagnosed (or future) samples can potentially be truly causal to the disease. Since our dataset is not an unbiased sample of MDM patient population, to interpret ANEVA-DOT findings we estimated positive predictive value (PPV) and false positive rates (FPR) of our analysis under two extreme scenarios. First, we estimated the fraction of ANEVA-DOT genes that are true positive (PPV) under the assumption that every patient has one regulatory disruptive pathogenic mutation with available AE data. To do this we used cases shown in **Fig S11**. We found that despite the high sensitivity and the great improvement in specificity, only 5.4% ANEVA-DOT genes are true positives when all genes are considered, underpinning the complexity of identifying novel genes in rare disease (**Fig 4H**). The PPV increased to 62% when we limited our analysis to only known MDM genes (**Fig 4H**). Next, we looked at the fraction of negative cases that can lead to false positives (defined as presence of at least one MDM associated ANEVA-DOT gene) under a worst-case scenario that none of the patients carry a regulatory disruptive pathogenic mutation. To do this we used cases demonstrated in **Fig S12A**, and **Fig S13**. We found a 27% FPR under this scenario. Thus, we expect under worst case scenario to have 8.8 out of the 12 patients with known MDM ANEVA DOT gene to be false positive, and under a best case scenario we would expect up to 10.5 of the 17 known MDM, and up to 18.8 of the 349 identified ANEVA-DOT genes to be true positives.

### 2.3 Supplementary text 3: Further analysis of previously undiagnosed cases in MDM cohort

In the 33 cases for which previous genetic diagnosis had not been reached by WES/WGS or previous RNA-seq analysis, ANEVA-DOT led to a new confirmed diagnosis in one case, prioritized strong candidates in six cases, and identified possible candidates in 11 cases. These cases are detailed below:

#### Case N10

In patient N10, with no prior strong candidates from WES, WGS or RNA-seq, ANEVA-DOT identified *DES* as the most significant gene with allele imbalance. The patient had a rare (gnomAD AC = 0) missense variant at chr2:220283753 T>C (p.Leu190Pro) in *DES*, which, the ANEVA-DOT outlier status, suggested a possible compound heterozygous effect that we further inspected in RNA-seq data. This resulted in the identification of pseudo-exon insertion of 116 bps. The patient carries a 2:220289644 G>A intronic variant at the junction, which results in the creation of an acceptor splice site as GG>AG. This results in the activation of a native intronic GT splice site downstream. Importantly, the missense variant is paternally inherited and the intronic variant is maternally inherited, and both variants are found in *trans* in the proband’s affected sibling. The intronic splice gain was missed previously by Cummings et al. due to the proportionally low read support for the pseudoexon (40 and 36 reads) compared to wild type splicing (8031 reads), highlighting the value of a prioritization framework based on allele imbalance at identifying new diagnoses that were previously missed.

#### Case N40

In patient N40, an adult patient with slowly progressive weakness, ANEVA-DOT identified *HADHA* as the top gene with allele imbalance. Disruption of HADHA results in autosomal recessive mitochondrial trifunctional protein deficiency, which manifests as generalized slow progressive weakness, offering a possible phenotype match. The patient carries a paternally inherited, rare missense variant with AC = 76 in gnomAD (no homozygotes). The haplotype carrying the heterozygous missense variant represents 100% alleles in RNA-seq data, indicating complete absence of the *trans* haplotype. The patient carries a rare intronic variant 2:26239296 C>T (AC = 2 in gnomAD) that we hypothesize may be resulting in branch point motif disruption. Given the segregation evidence, the phenotype match and the allele imbalance, this gene is now being pursued as a strong candidate for the sample, and functional studies to assess splicing around the intronic variant are in progress, again highlighting the value of gene prioritization.

#### Cases N26 and E1

In patients N26 and E1, both with clinically diagnosed nemaline myopathy, ANEVA-DOT highly prioritized *NEB*. Patient N26 carries a missense variant in *trans* with a frameshift variant in the NEB triplicated region. Functional analyses are underway to evaluate any possible splice defects caused by the missense variant. Similarly, patient E1 and their affected sibling carry a stop gained variants in NEB, which is *in trans* with several intronic variants. Further variant interpretation is required to identify the causative variant in the gene for the patients well-fitting phenotype.

#### Case N38

For patient N38 with multiple congenital abnormalities include congenital myopathy, respiratory insufficiency and cleft palate, ANEVA-DOT prioritized *GBE1, ATP2A1*, and *TNNT3*. The patient carries a missense and nonsense variant in trans in GBE1, which is characterized as an incidental finding, and a rare non-coding variant in *ATP2A1* marked for follow up.

#### Cases N24 and N23

In patient N24 with axial myopathy and respiratory insufficiency, ANEVA-DOT prioritized *COL6A1*. The patient carries a frameshift variant in the gene, but no second hit. Similarly, in patient N23 with congenital myopathy, ANEVA-DOT prioritized *COL6A3* which carries an essential splice variant shared by the affected sibling, but no second hit.

Overall, the strong candidates provided by ANEVA-DOT for follow-up helped prioritize genes and variants, but it is important to note that gene prioritization, even in cases with complete allele imbalance, still requires careful variant interpretation and functional assessment to identify the causative variant to complete a diagnosis. In addition to these stronger candidates in known muscle disease gene, ANEVA-DOT identified possible candidates in 11 cases that are currently under assessment for diagnosis.

### 3 Supplementary table legends

**Table S1: V^G^ estimates from AE data**. AE-based estimates of V^G^ for all tissues in GTEx data, and for GEUVADIS data, as well as average V^G^ across GTEx tissues, 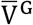, for all available gene.

**Table S2: V^G^ estimates from eQTL data**. eQTL-based estimates of V^G^ for all tissues in GTEx data, as well as average V^G^ across GTEx tissues, 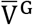, for all available gene.

**Table S3: Gensets used in figure 3D**.

**Table S4: Blacklisted genes in ANEVA-DOT**. List of false positive prone genes that were excluded from ANEVA-DOT analysis of GTEx muscle and MDM data due to their high rate of occurrence as dosage outlier in GTEx data. Additionally, the file includes lower bound for occurrence of ANEVA-DOT outliers in all available genes and tissues in GTEx data.

**Table S5: MDM analysis details**. Sample-level information for all patients included in the MDM cohort, including phenotypes, previous diagnosis, ANEVA-DOT results, and downstream analysis verdict. Additionally, the file includes the list of MDM associated genes used in the analysis.

### 4 Supplementary figures

**Fig S1:**
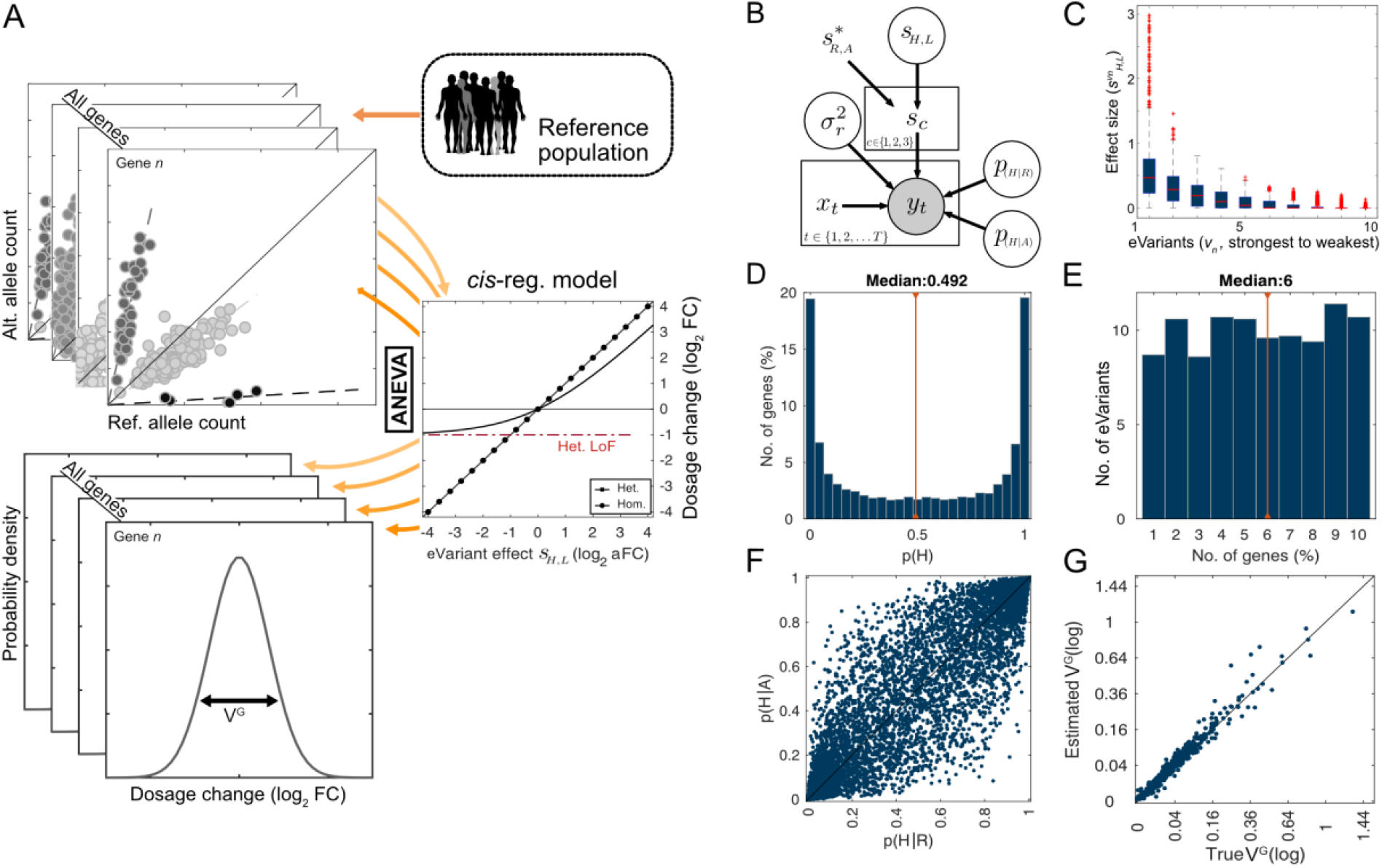
The ANEVA model and simulations. A) Schematic representation of ANEVA. ANEVA uses a generative model of population AE data and a mechanistic model *cis*-regulatory variation to estimates the magnitude of genetic variation in expression consistently across genes. B) Bayesian plate diagram of the population AE model used for ANEVA. See **Eq. 20–22** for description of variables and their dependency structure. C-F) Simulation parameters with the effect size distribution of regulatory variants (C), the frequency of the higher expressed regulatory allele of the main variant (D), number of regulatory variants per gene (E), and linkage disequilibrium between aeSNP and the main regulatory variant (F). G) Simulation results for V^G^. The estimated V^G^ values using ANEVA (y-axis) are consistent with true V^G^ values used for simulation of AE data (x-axis).

**Fig S2:**
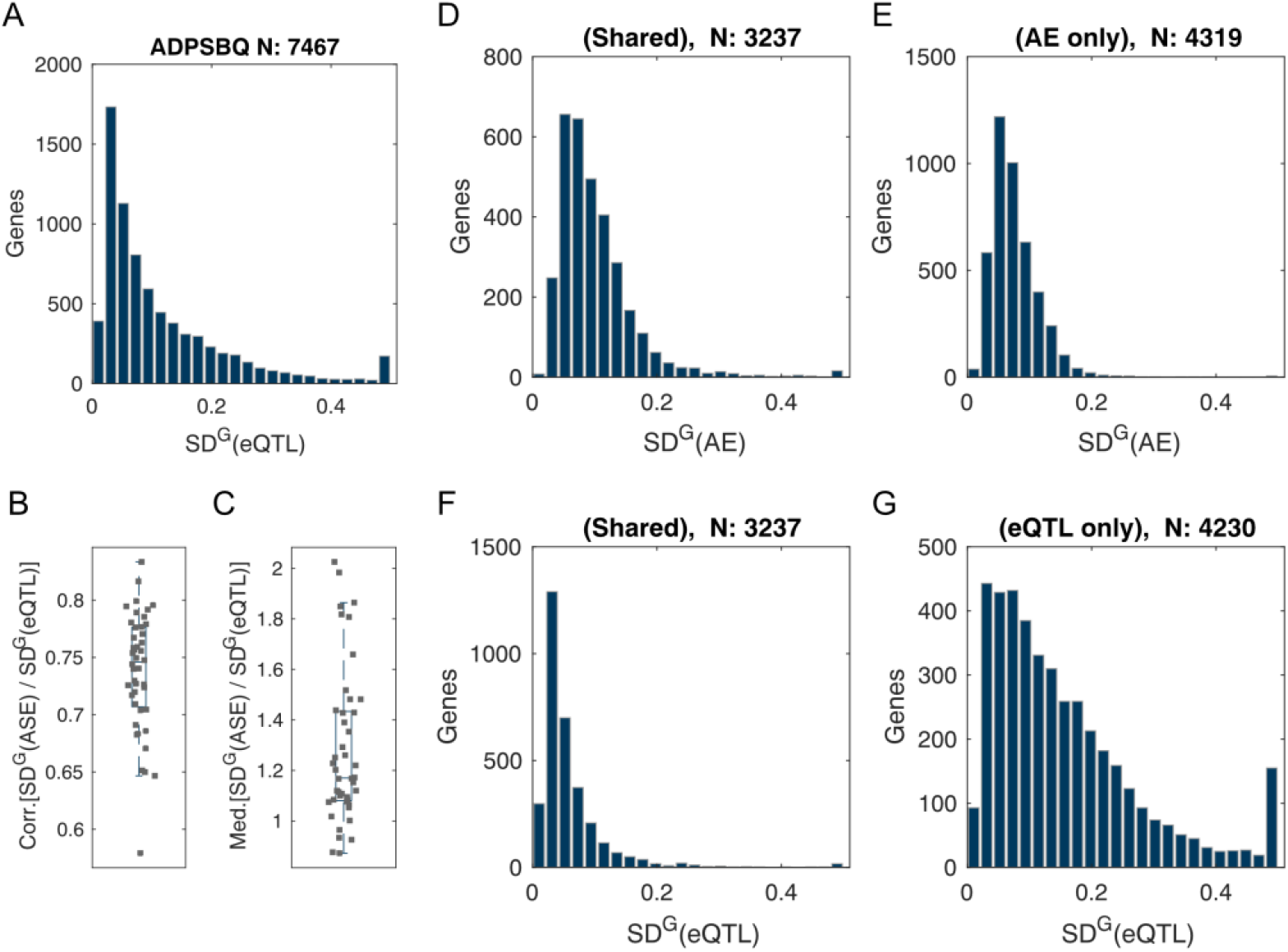
Comparison of ANEVA estimates from AE and eQTL data. A) SD^G^ (square root of V^G^) estimates for GTEx adipose subcutaneous calculated from eQTL data. B) correlation and C) median ratio of the AE based estimates and eQTL based estimates for each tissue. Estimates from AE and eQTL data were consistent (median rank correlation 0.75), but AE-based estimates were systematically larger for genes available in both (median ratio 1.17), possibly because AE data unlike eQTLs capture the effect of all regulatory variants including those that are rare. D-E) SD^G^ distribution calculated from AE data for genes that have both AE and eQTL estimates (D), and for genes with AE estimates only (E). F-G) SD^G^ distribution calculated from eQTL data for genes that have both AE and eQTL estimates (F), and for genes with eQTL estimates only (G). Altogether, these figures and **Figure 2B-C** show that the eQTL and AE ANEVA analyses yield similar estimates, but each tend to be available for very different types of genes.

**Fig S3:**
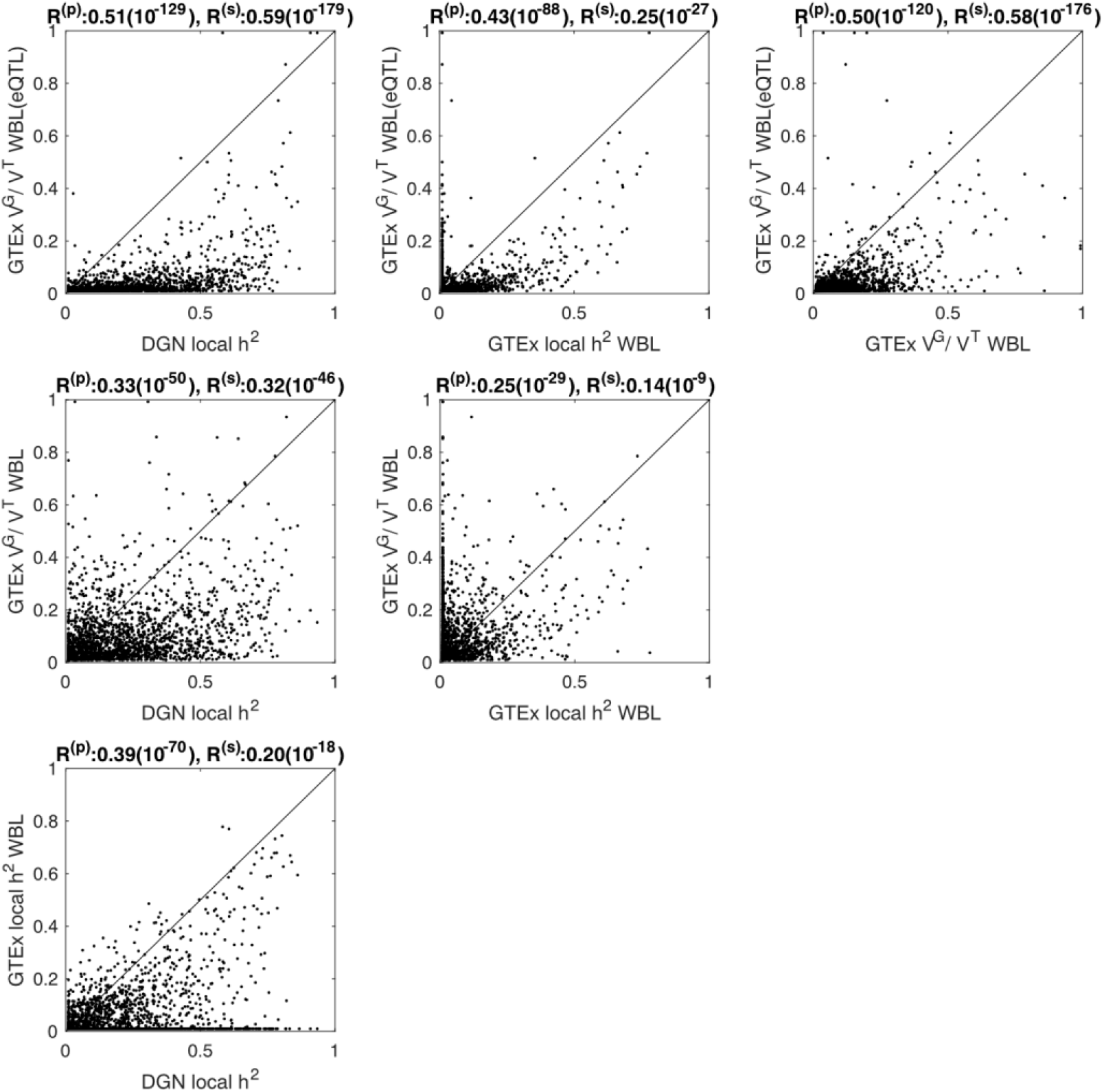
Comparisons of local heritability (h^2^) estimates. Scatter plots of all h^2^ comparisons for 1939 whole blood genes available in all four datasets: DGN data set local h^2^ (from BSLMM), GTEx whole blood (WBL) local h^2^ (from BSLMM), and ANEVA’s V^G^/V^T^ calculated from GTEx whole blood using either AE or eQTL data. The correlations are noisy overall, which is unsurprising for relatively low sample sizes. Additionally, the ANEVA estimates are systematically smaller. This is due to the fact that we did not correct for any confounding factors in the expression data when calculating V^T^, whereas the BSLMM models (similarly to standard *cis*-eQTL analyses) included PEER factors to remove technical and other transcriptome-wide variation in gene expression. We did not include PEER factors to make sure the observed correlation between our estimates and BSLMM are completely independent and are not inflated by this step.

**Fig S4:**
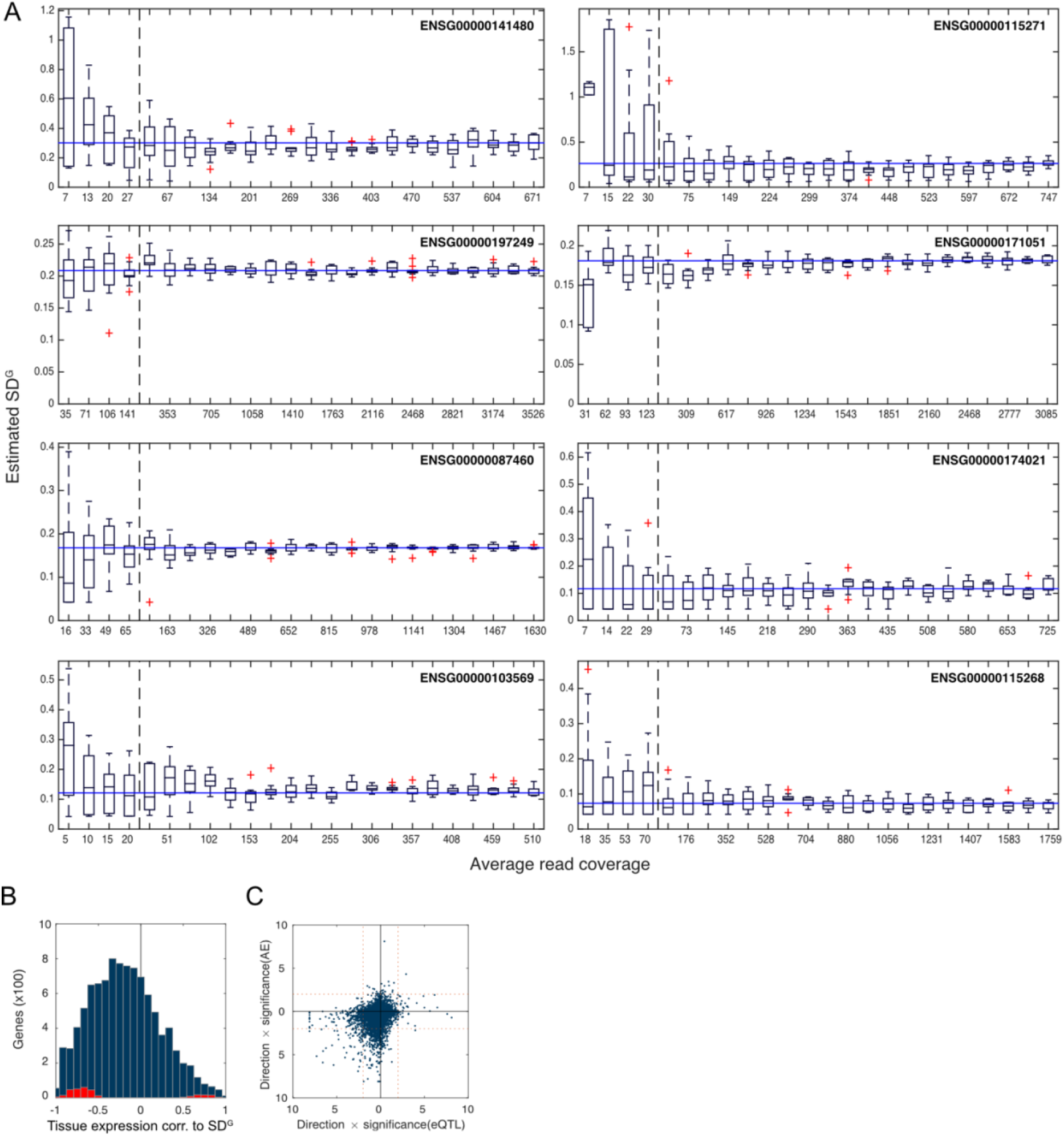
V^G^ estimates across tissues. A) ANEVA estimates from down-sampled data. Shown are SD^G^ estimates as a function of average read count for 768 down-sampling experiments including eight genes, 24 down-sampling rates, and four independently drawn samples at each rate. Variance of the estimates grows as the read coverage shrinks but no consistent monotonic pattern is observed. The solid blue line shows the SD^G^ estimate from the original data, and each box shows the estimate from the four samples at that rate. To the left of the dashed line coverage rate increases at 1% intervals, and to the right at 5%. B) Distribution of rank correlation values between eQTL-based estimates of V^G^ in a given tissue and the expression of gene in the tissue for all available genes. Genes individually showing significant correlation are shown in red (5% FDR). C) Observed patterns of V^G^-expression correlation are consistent between eQTL-based and AE-based estimates. Shown are 5554 genes with V^G^ estimates in at least five tissues from both AE and eQTL data. Each dot is a gene and the coordinates are *Sign*(R^(s)^) × −log_10_ *p*, where R^(s)^ is the rank correlation between V^G^ and expression level across tissues for the gene, and *p* is the p-value associated with R^(s)^. Values derived using eQTL-based estimates are on x-axis, and those from AE-based estimates are on y-axis. The generally consistent results confirms that the correlation between V^G^ and expression is not driven by AE-based results being biased by low read counts.

**Fig S5:**
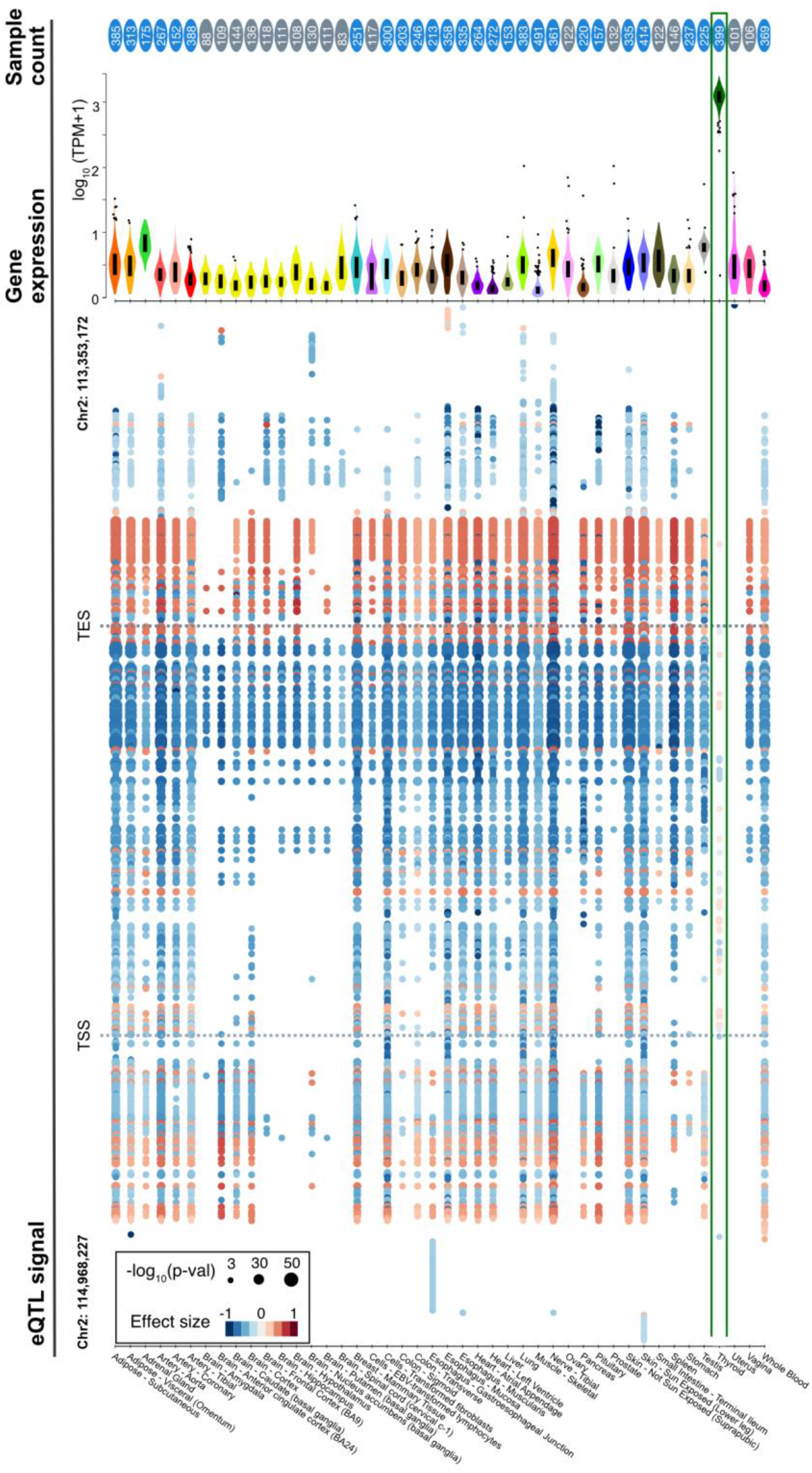
Tissue-specific expression and regulatory variation in *PAX8*. The *PAX8* gene has a weak eQTL signal in thyroid tissue despite high expression and large sample size. The figure shows all 6738 eVariants for PAX8 in GTEx v7 data without LD pruning. Details of all eQTL signals for PAX8 are available for browsing at: https://gtexportal.org/home/gene/PAX8

**Fig S6:**
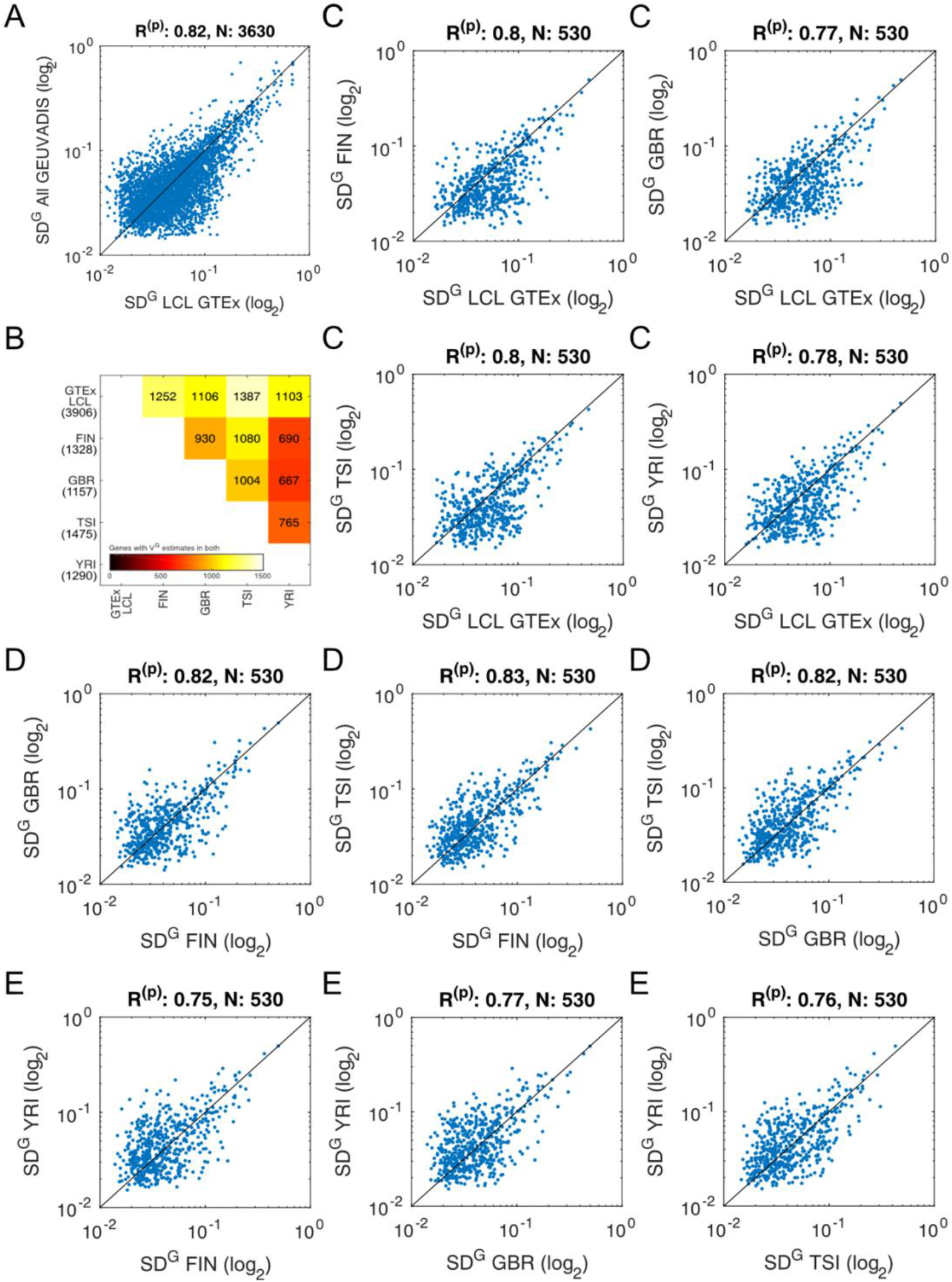
ANEVA estimates from different populations. A) ANEVA estimates of SD^G^ (square root of V^G^) in lymphoblastoid cell lines (LCL) calculated from GTEx and GEUVADIS datasets, where R^(p)^ is the Pearson correlation, and N is the number of genes plotted. B) Number of genes with V^G^ estimates available in each possible pair of GEUVADIS subpopulations and GTEx LCLs. Number in parenthesis are the total number of genes available in each subpopulation. C) SD^G^ estimates from GTEx compared to estimates from each subpopulation in GEUVADIS for genes available in all datasets. D-E) GEUVADIS European subpopulations compared to each other, and to the Yoruba (YRI) population (E) for genes available in all datasets. All in all, the genes available for V^G^ calculation in African and European sub-populations are different due to population-specificity of common aeSNPs. However, when AE data is available in both groups, estimates of genetic variation in expression from European sub-populations are not drastically more correlated to each other than to those from Yoruba population.

**Fig S7:**
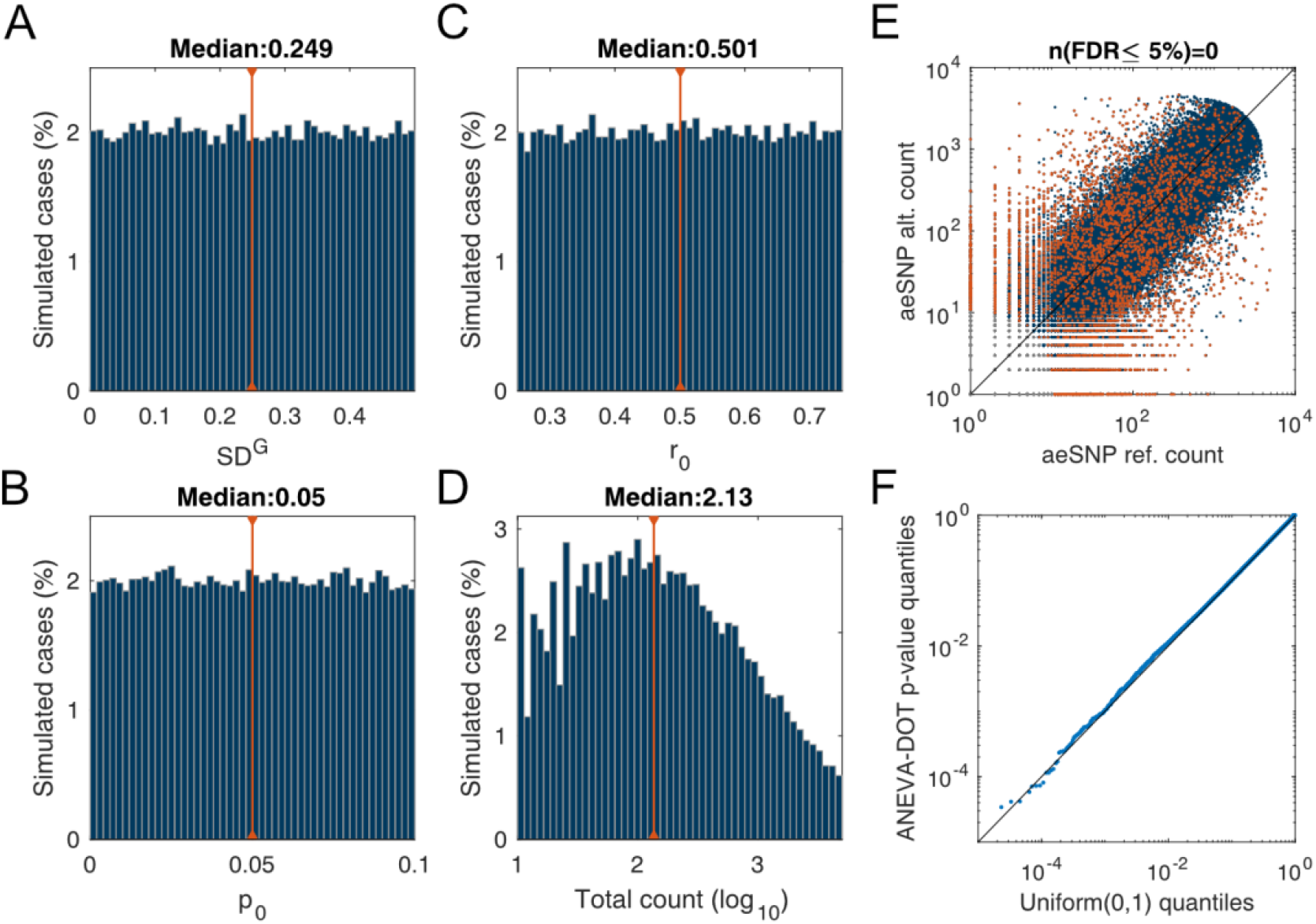
ANEVA-DOT simulation study. We generated 100,000 simulated AE samples for the null model. Each sample was generated and tested using a unique set of randomly generated parameters to ensure ANEVA-DOT is well calibrated across all parameter combinations. A-D) Distribution of simulated parameters: square root of V^G^ (SD^G^), reference bias (*r_0_*), sequencing noise (*p_0_*), and read coverage (total count) used for generating AE data. Red line indicates the median value. E) Simulated AE data and ANEVA-DOT results. Red dots are cases with nominal p-value ≤ 0.05. These are dispersed over all allelic imbalance values due to the fact that each point is tested against its own null model defined by SD^G^, *r_0_*, and *p_0_* in addition to the read coverage. None of the simulated data points were significant at 5% FDR. F) QQ-plot showing ANEVA-DOT p-values are consistent with theoretical expected distribution and thus calibrated.

**Fig S8:**
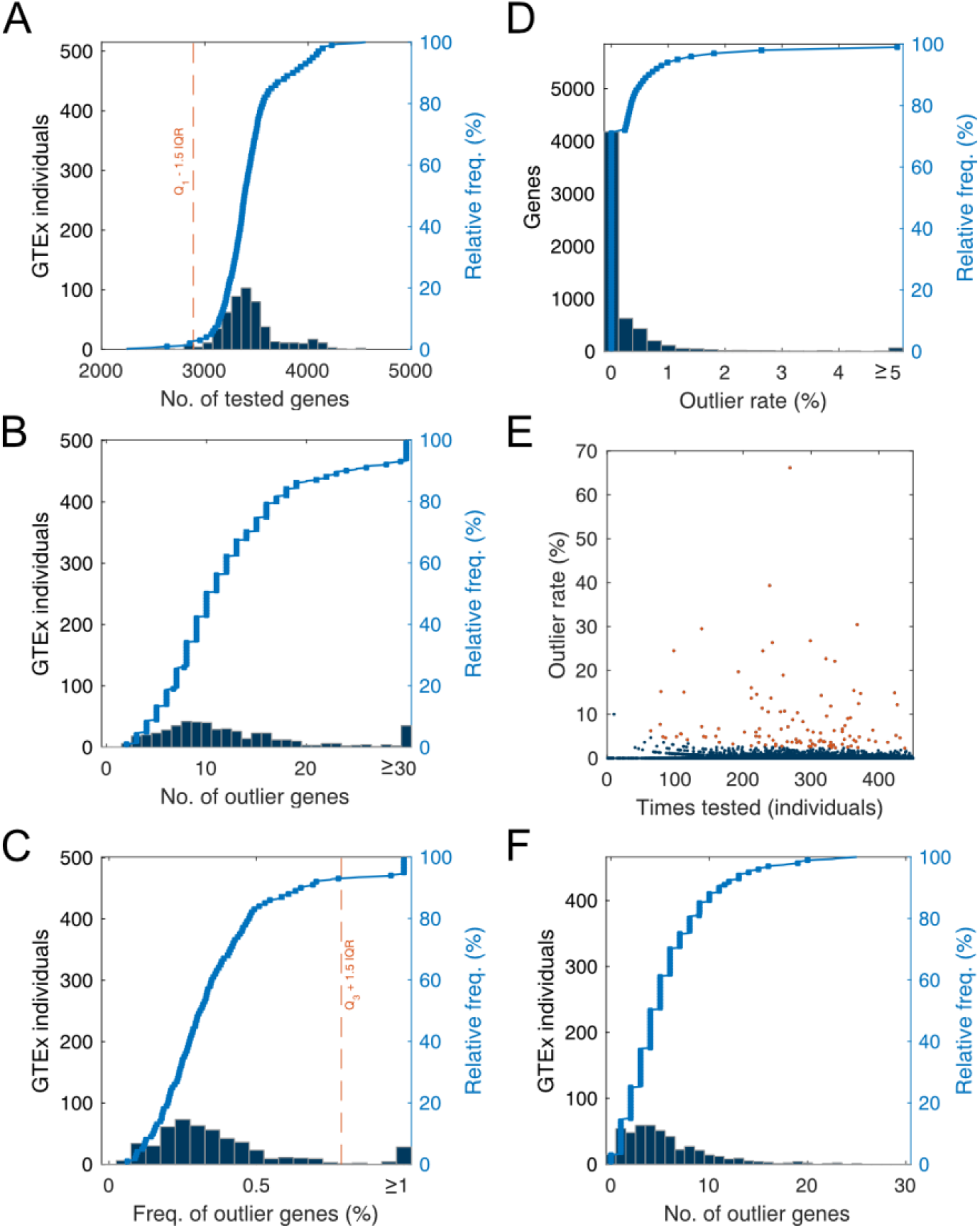
ANEVA-DOT in GTEx. Summary statistics from muscle tissue with A) number of tested genes per individual, B) the number of outlier genes per individual, C), the percentage of outlier genes per individual, and D) the percentage of outlier individuals per gene. E) Genes excluded from downstream analysis due to too many outliers in them. F) The final number of ANEVA-DOT genes per individual across the 466 high quality GTEx muscle samples passing quality controls. In total, we found a total of 2554 cases of gene-individual outlier pairs.

**Fig S9:**
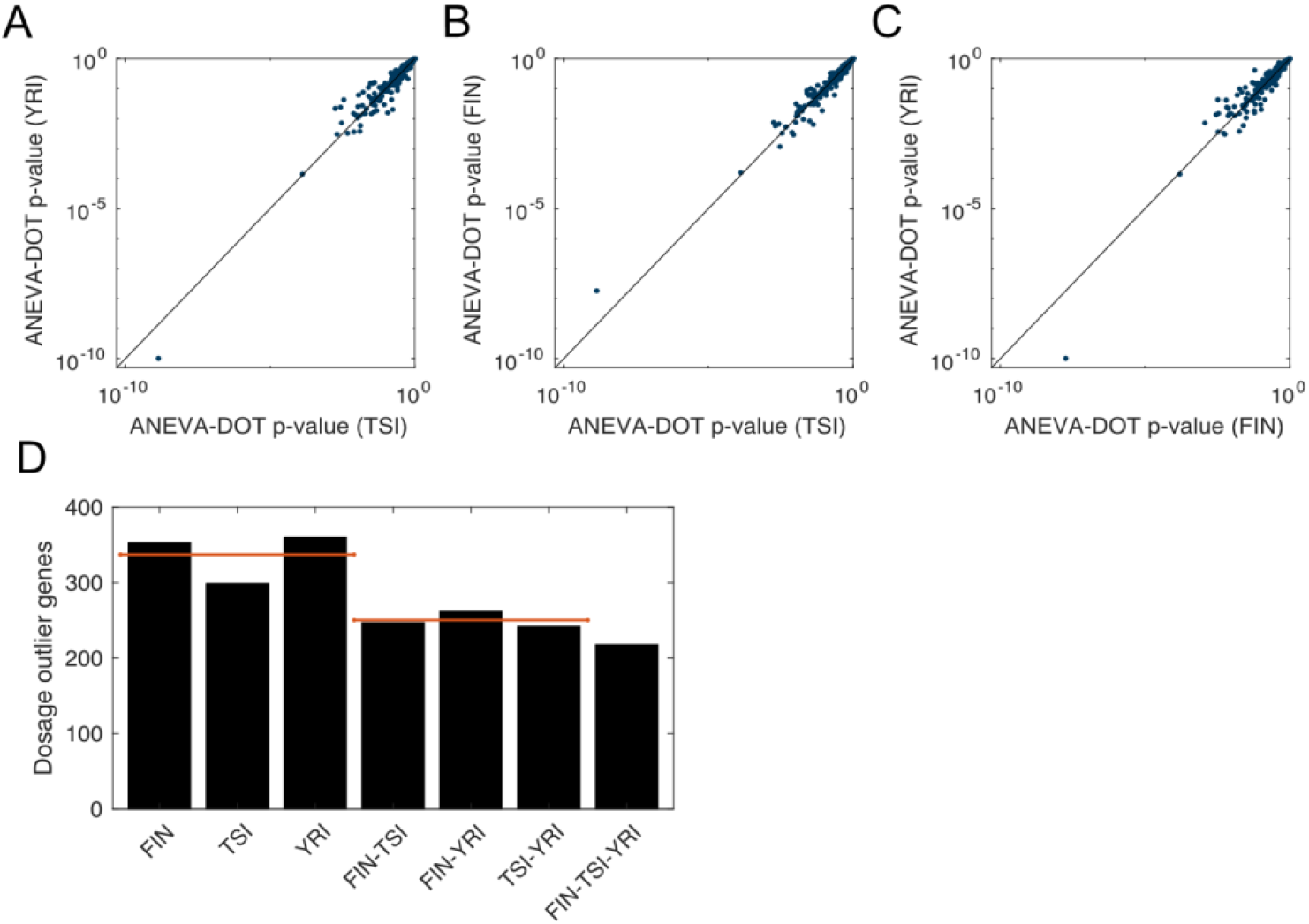
ANEVA-DOT across populations. We estimated ANEVA-DOT outliers for GEUVADIS samples from the GBR (British) population using V^G^ reference estimates from other populations. A-C) direct comparison of p-values. D) Intersection of 5% FDR genes for the entire GBR sample set across different reference sub-populations. Red lines indicate the median of the three bars.

**Fig S10:**
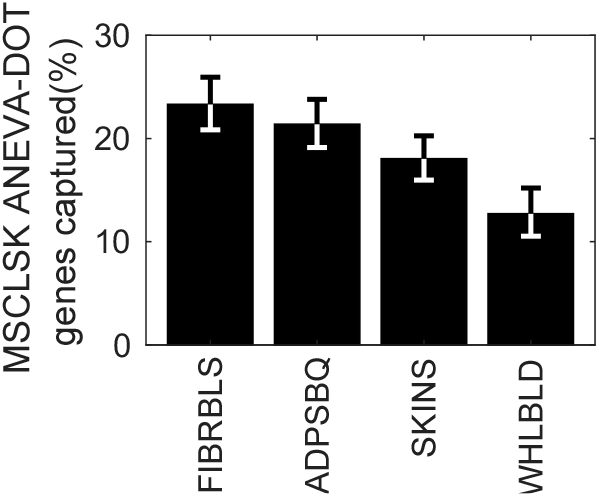
ANEVA-DOT across tissues. ANEVA-DOT applied to GTEx muscle samples but using V^G^ estimates from three other tissues as the population reference. The y-axis shows the percentage of ANEVA-DOT outlier genes the muscle reference that are identified with the different tissue references (5% FDR).

**Fig S11:**
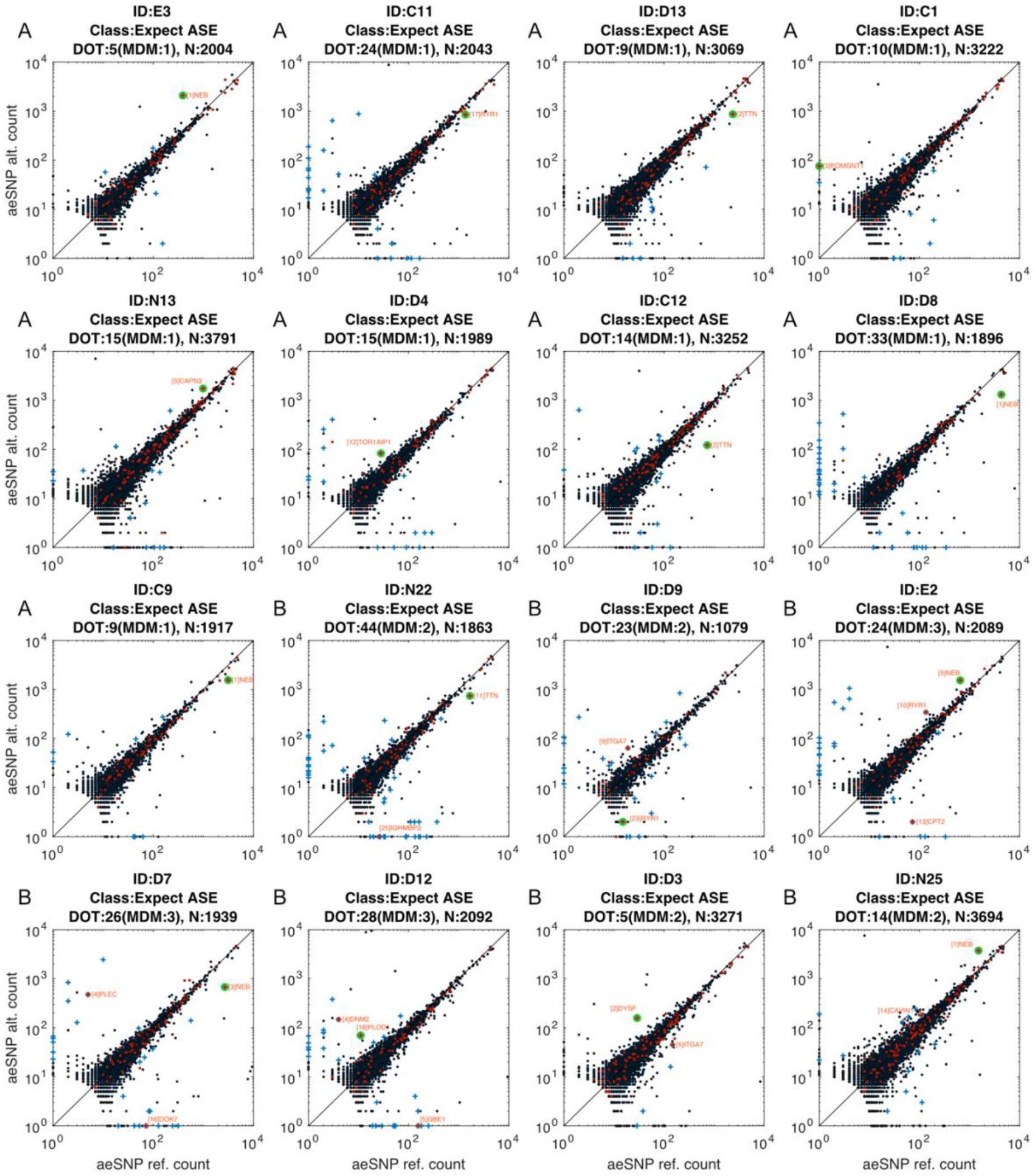
MDM patients where ANEVA-DOT corroborates the previous diagnosis. AE samples from patients with a previous molecular diagnosis. Based on the nature of the underlying pathogenic mutation the affected gene is expected to show excessive allelic imbalance (designated by green circle). Here, we show individuals where the diagnosed gene is among the ANEVA-DOT outliers. The individuals in the (A) panels have only one MD ANEVA-DOT gene, whereas the (B) individuals have several. Red: MDM gene, Blue: ANEVA-DOT gene. Known MDM ANEVA-DOT genes are named and numbered by their p-value rank in the entire sample. On the top the numbers are the number of DOT genes, the number of MDM DOT genes, and the total number of tested genes.

**Fig S12:**
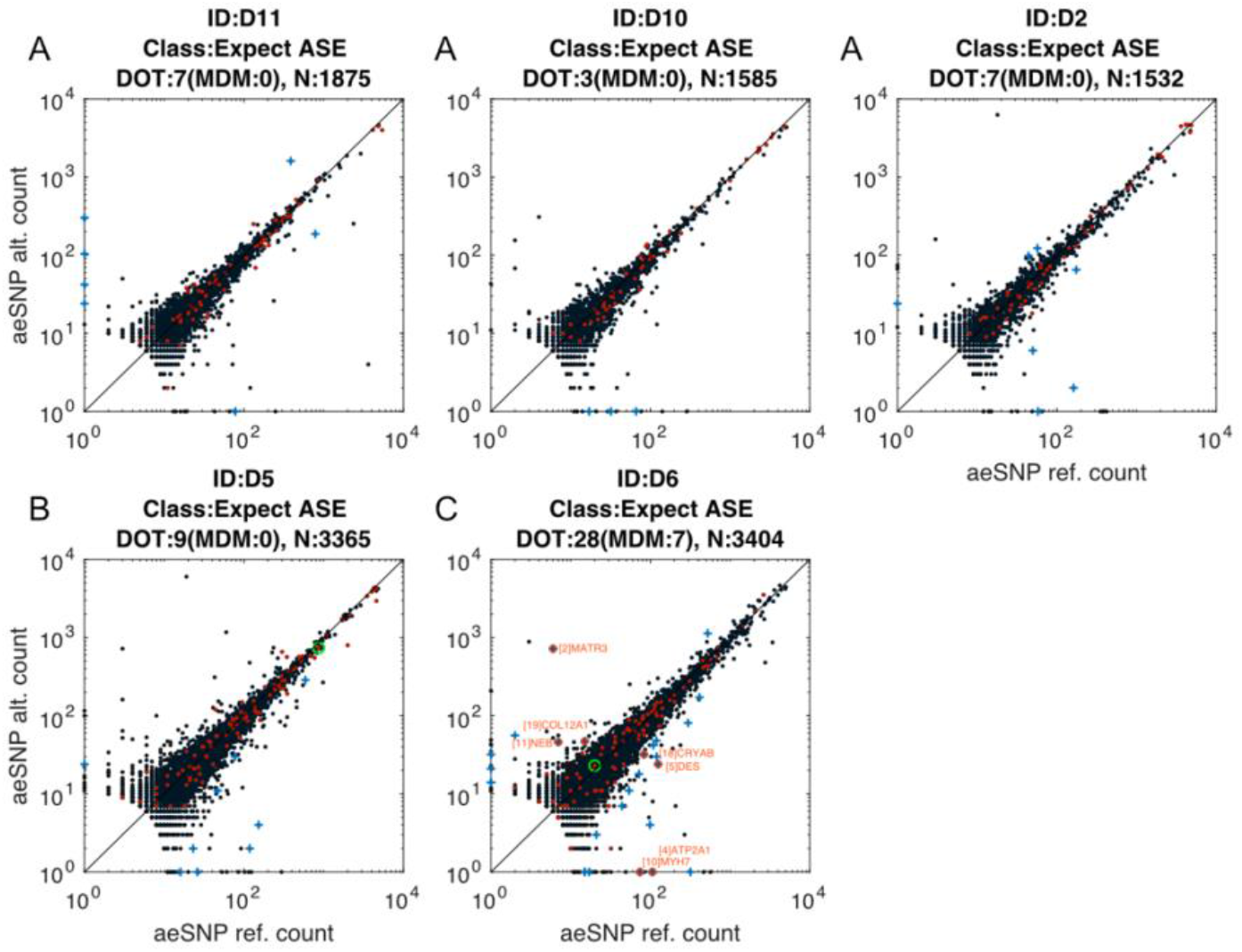
ANEVA-DOT results on MDM patients where the previous diagnosis is not corroborated. AE samples from patients with a previous molecular diagnosis. Based on the nature of the underlying pathogenic mutation the affected gene is expected to show excessive allelic imbalance (designated by green circle when available). However, these genes are not ANEVA-DOT outliers. A) panels are cases where the affected gene is entirely missing from AE data due to lack of aeSNP or minimal expression. These samples were included for estimating the empirical recall rate (as positive controls), and false discovery rate calculation (as negative controls) but were excluded from idealized recall and positive predictive value calculation where we aimed to find an upper bound under an ideal scenario. The individuals in (B, C) panels are cases where the affected gene does not have allelic imbalance under the standard binomial test and thus, statistically does not carry any signal for allelic imbalance. These samples were included for estimating the empirical recall rate (as positive controls) but were excluded from idealized recall and positive predictive value calculation where we aimed to find an upper bound under an ideal scenario. We note that the C) individual is an ANEVA-DOT outlier for multiple MDM genes, which might be additional or alternative disease-relevant findings to complement the first diagnosis. Red: MDM gene, Blue: ANEVA-DOT gene. Known MDM ANEVA-DOT genes are named and numbered by their p-value rank in the entire sample. On the top the numbers are the number of DOT genes, the number of MDM DOT genes, and the total number of tested genes.

**Fig S13:**
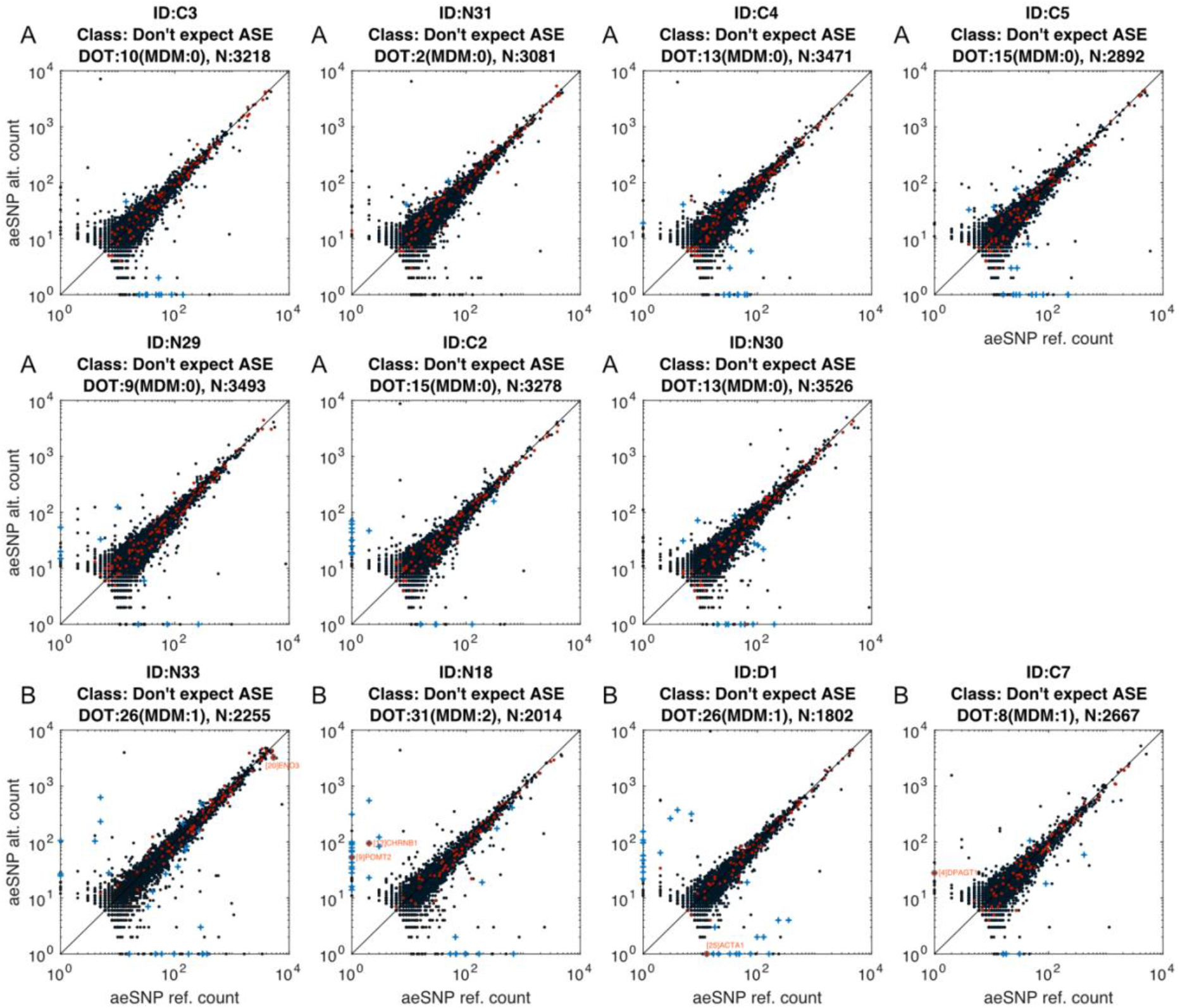
ANEVA-DOT results on MDM patients where no AE is expected. AE samples from patients with previous molecular diagnosis. Based on the nature of the underlying pathogenic mutation the affected gene is either not present in AE data (e.g. in case of male patients with Duchenne muscular dystrophy) or is not expected to show allelic imbalance. These samples represent “negative controls” and were included for false discovery rate calculation (along with samples in **Fig S12A**) where we look at false discovery rate under a worst case scenario when every sample is negative. The individuals in (A) panels had no MDM ANEVA-DOT genes, and those in (B) panels had MDM ANEVA-DOT genes. Red: MDM gene, Blue: ANEVA-DOT gene. Known MDM ANEVA-DOT genes are named and numbered by their p-value rank in the entire sample. On the top the numbers are the number of DOT genes, the number of MDM DOT genes, and the total number of tested genes.

**Fig S14:**
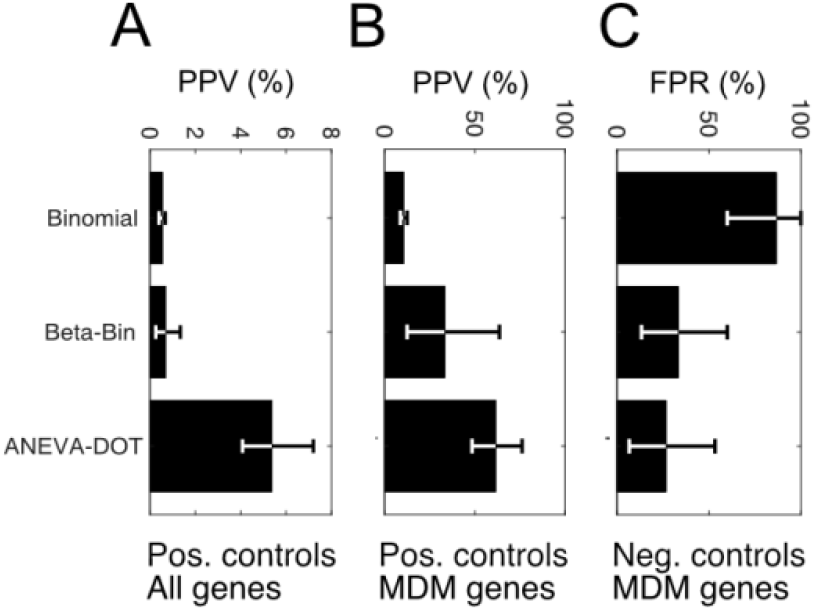
Positive predictive and false discovery rates for AE tests. A-B) Positive predictive value (PPV) of ANEVA-DOT genes that is the fraction of ANEVA-DOT genes truly causal to the disease when all genes (A), and when only known MDM genes (B) are considered. PPV is calculated under the best case scenario where, 1) the disease-causing mutation leads to allelic imbalance and 2) it is available in AE data in every patient. Each patient has only one disease causing mutation. C) False positive rate (FPR) calculated as the fraction of patients with at least one ANEVA-DOT gene that is a known MDM gene under worst case scenario when none of the patients carry a mutation causing allelic imbalance that is available in AE data (i.e. when any ANEVA-DOT gene is a false positive).

**Fig S15:**
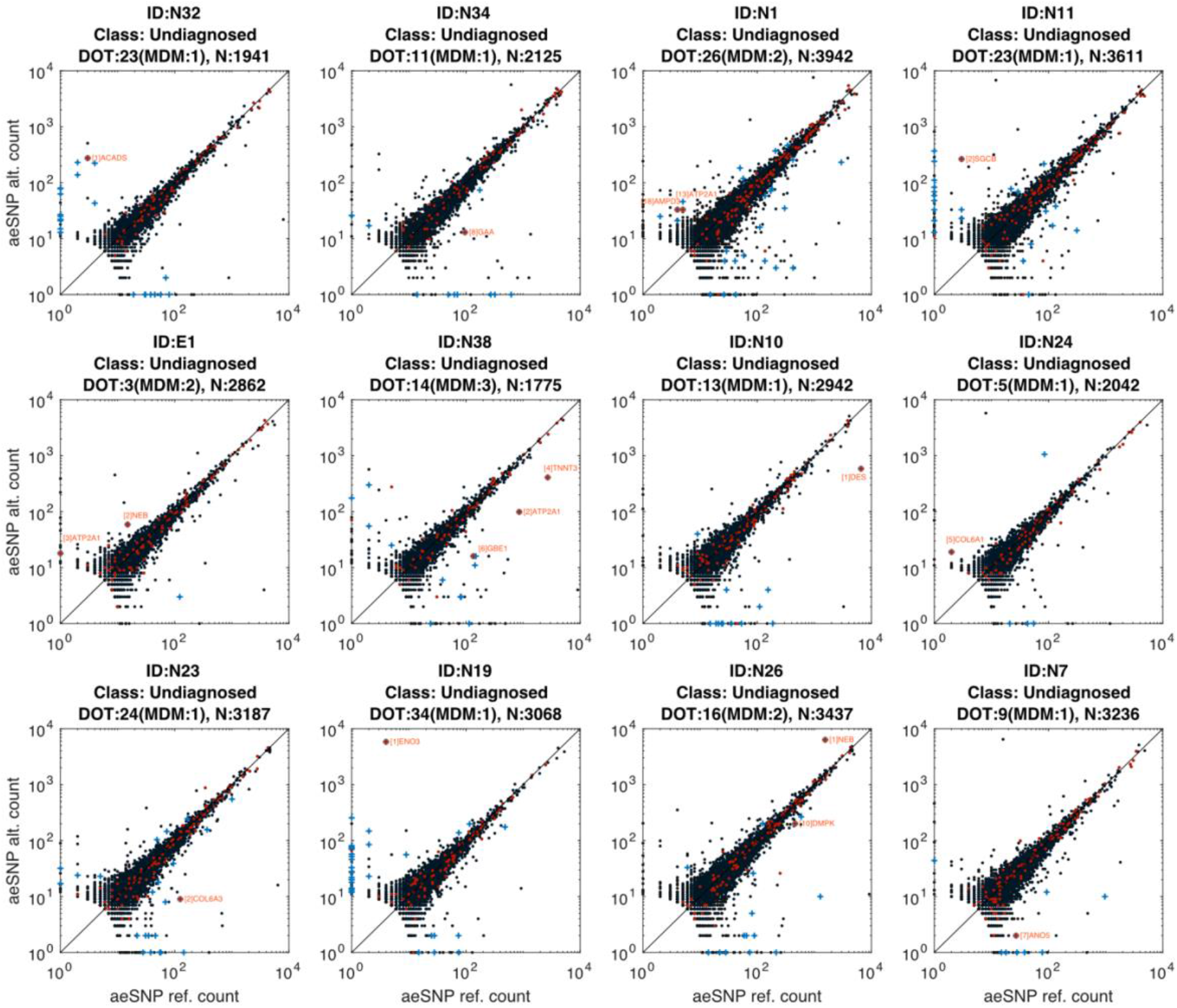
ANEVA-DOT results on undiagnosed MDM patients where our newly identified candidates include known MDM genes. AE samples from patients without previous molecular diagnosis. In these samples we find one or more known MDM genes to be an ANEVA-DOT outlier. Red: MDM gene, Blue: ANEVA-DOT gene. Known MDM ANEVA-DOT genes are named and numbered by their p-value rank in the entire sample. On the top the numbers are the number of DOT genes, the number of MDM DOT genes, and the total number of tested genes.

**Fig S16:**
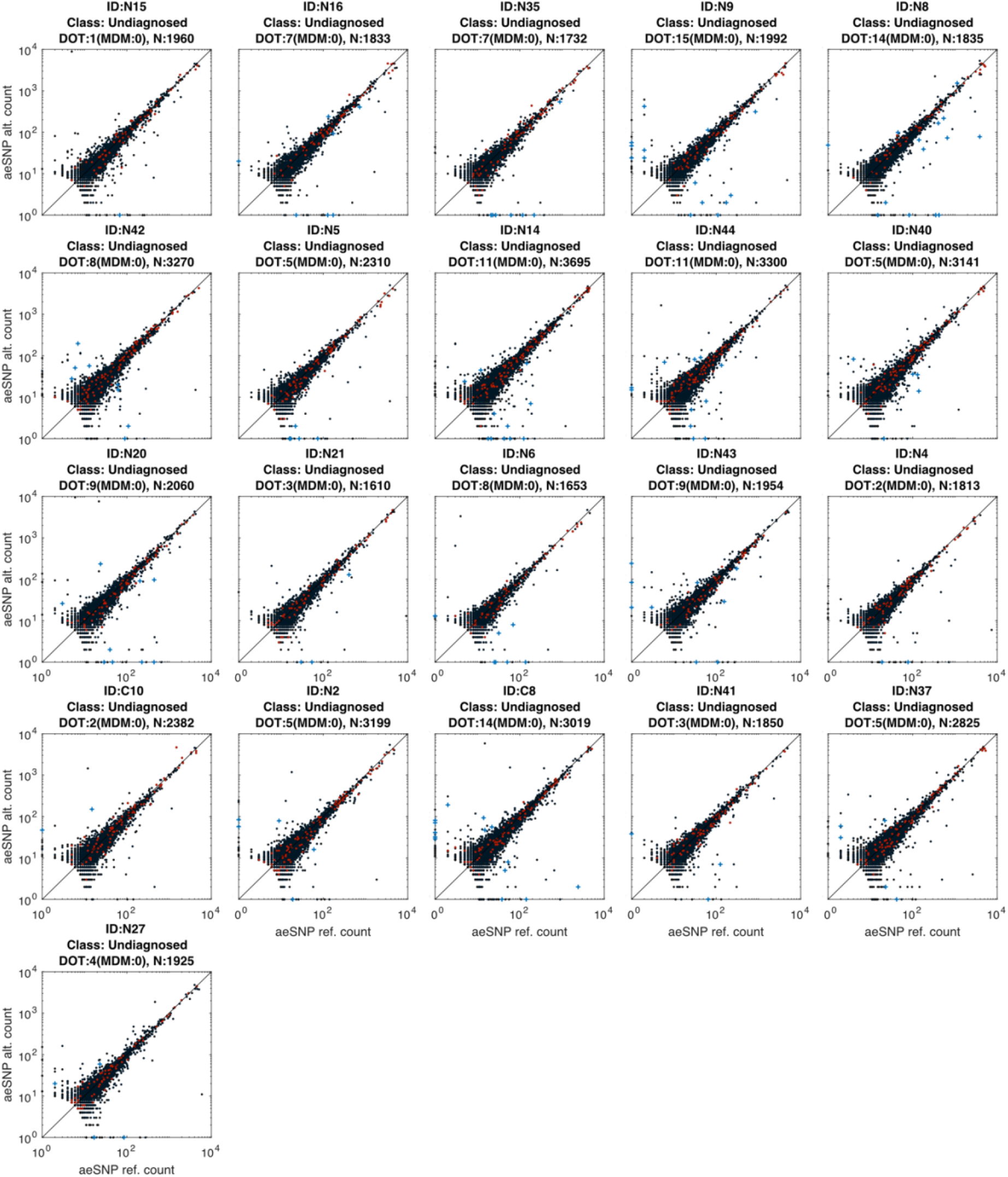
ANEVA-DOT results on undiagnosed MDM patients where our newly identified candidates do not include any previously known MDM genes. AE samples from patients without previous molecular diagnosis. In these samples we do not find any known MDM genes to be an ANEVA-DOT outlier. Red: MDM gene, Blue: ANEVA-DOT gene. On the top the numbers are the number of DOT genes, the number of MDM DOT genes, and the total number of tested genes.

**Fig S17:**
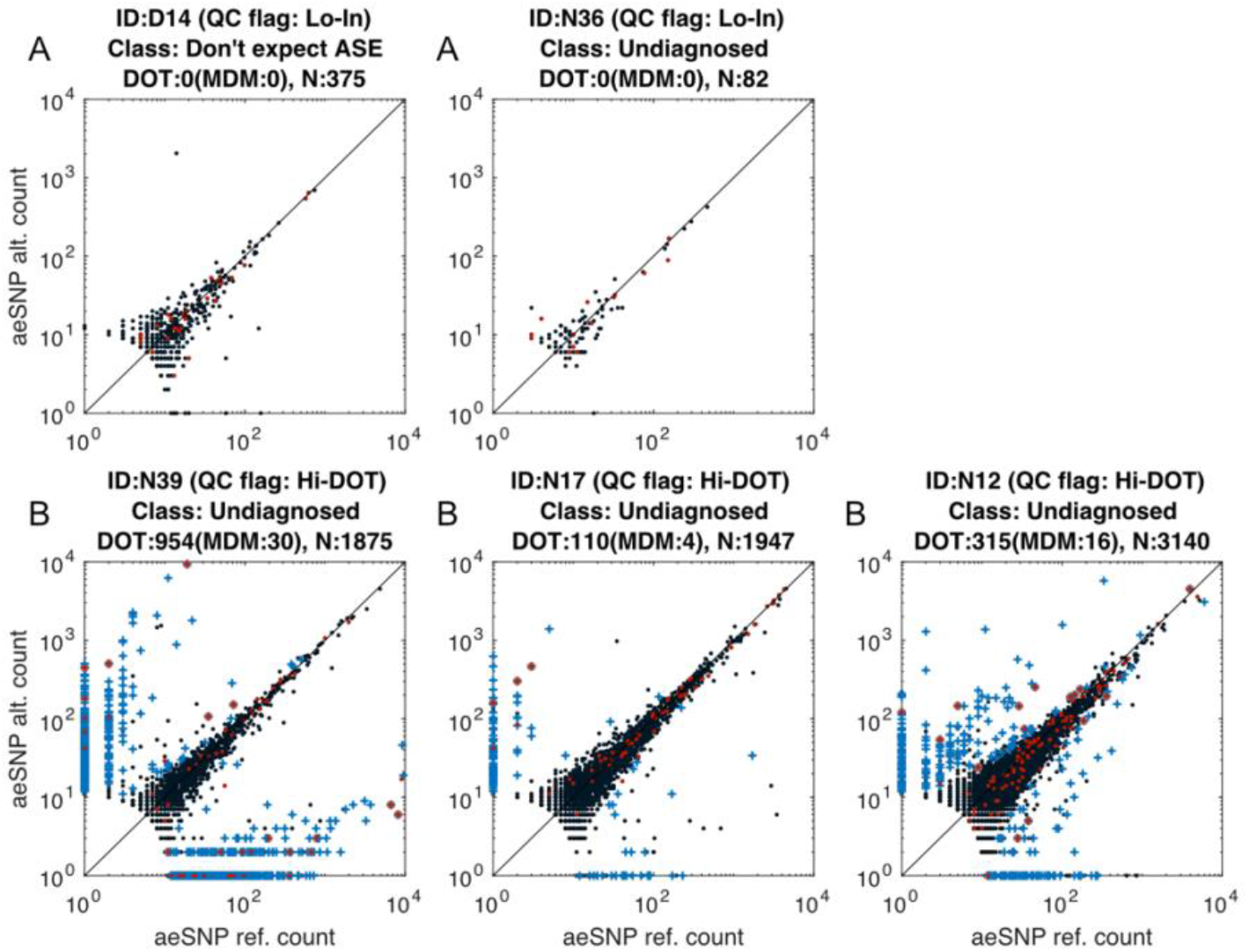
ANEVA-DOT results on MDM patients excluded from analysis. AE samples from these patients did not pass our quality control criteria. A) Too few genes were available in the data due to low coverage. We used Q_1_-1.5IQR from all tested samples as the threshold. B) Too many genes appeared as ANEVA-DOT outlier possibly due to genotyping error, sample contamination, or extensive PCR duplication that is not captured by duplicate removal. The threshold was set as Q_3_+1.5IQR from all samples except the two in panel A that did not pass the coverage threshold. Red: MDM gene, Blue: ANEVA-DOT gene. On the top the numbers are the number of DOT genes, the number of MDM DOT genes, and the total number of tested genes.

**Fig S18:**
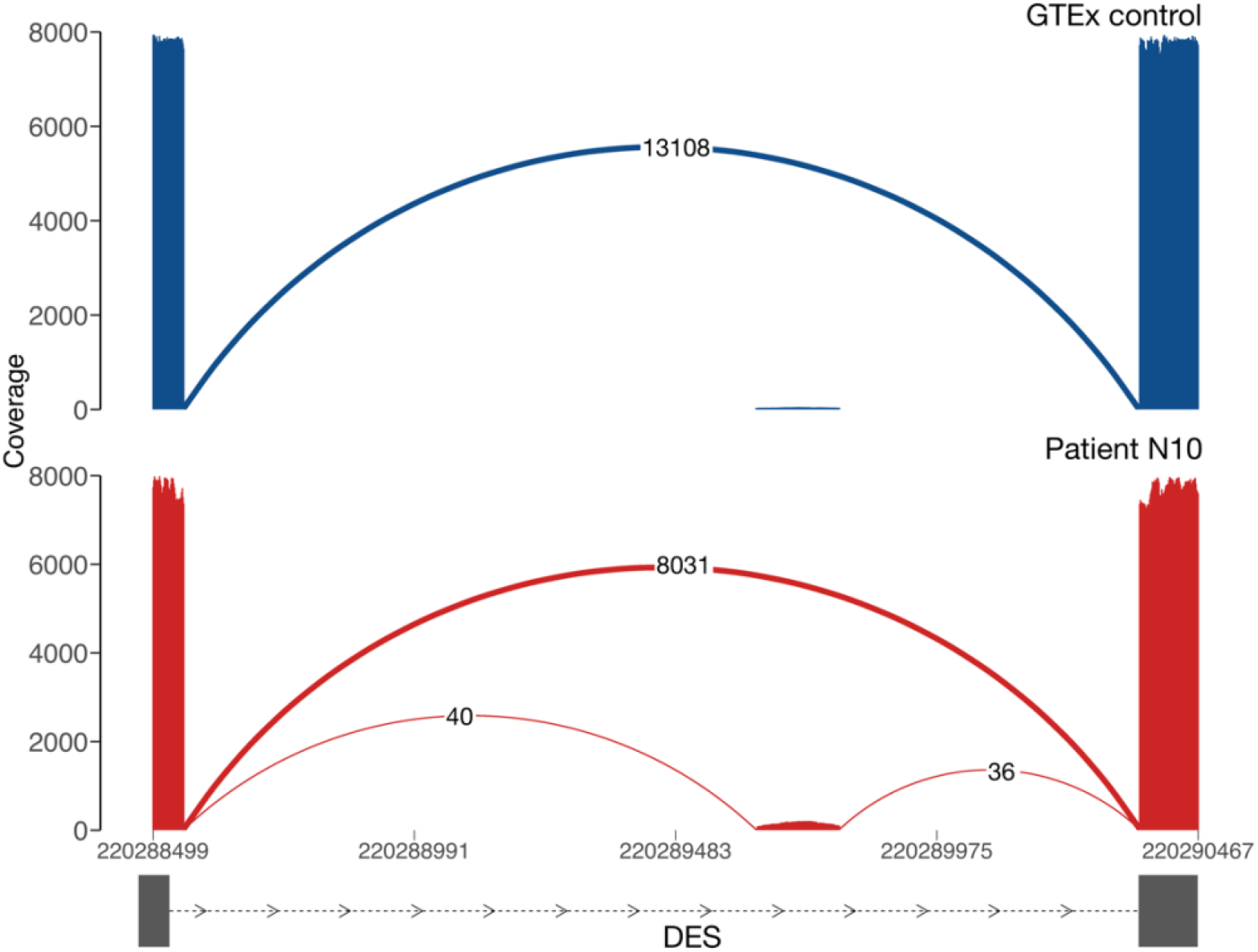
Sashimi plot showing pseudoexon insertion in patient N10. Patient N10 carries a heterozygous variant chr2:220289644 (c.1289-741G>A) resulting in the creation of a splice acceptor site, resulting in the inclusion of 116 out-of-frame bases into the transcript (see supplementary text 3 for details). Numbers in arches represent the number of uniquely mapping, non-duplicate, primary alignments that support the splicing event. The control sample in blue is GTEx skeletal muscle sample G18069.

